# *Plagl2* unlocks the latent regenerative potential of Müller glia in the adult mouse retina

**DOI:** 10.64898/2026.04.18.719349

**Authors:** Taimu Masaki, Mikiya Watanabe, Michiko Mandai, Miho Kihara, Takaya Abe, Ryoichiro Kageyama

**Affiliations:** RIKEN Center for Biosystems Dynamics Research, Hyogo 650-0047, Japan; Graduate School of Medicine, Kyoto University, Kyoto 606-8501, Japan; Research Center, Kobe City Eye Hospital, Hyogo 650-0047, Japan; Cell and Gene Therapy in Ophthalmology Laboratory, RIKEN BZP, Saitama 351-0198, Japan; Laboratory for Animal Resources and Genetic Engineering, RIKEN, Hyogo 650-0047, Japan; Research Organization of Science and Technology, Ritsumeikan University, Shiga 525-8577, Japan

**Keywords:** Müller glia, *Plagl2*, adult mouse retina, regeneration, neurogenesis, single-cell RNA sequencing

## Abstract

Müller glia (MG) in the adult mammalian retina have long been recognized as a potential endogenous source for the regeneration of lost retinal neurons. However, existing MG reprogramming strategies yield an incomplete regenerative response by favoring either proliferation or neurogenic differentiation. Here, we show that the zinc-finger transcription factor *Plagl2*, which has been demonstrated to rejuvenate aged neural stem cells, was sufficient to reprogram adult mouse MG into a progenitor-like state with coupled proliferative and neurogenic competence. Using histology, time-lapse imaging, and single-cell transcriptomics, we found that *Plagl2* drove regulated rounds of MG cell cycle re-entry, while N-methyl-D-aspartate-induced retinal injury further promoted the acquisition of neurogenic competence toward inner nuclear layer neuronal identities. These findings identify *Plagl2* as a novel rejuvenator of mammalian MG and support the general principle that reprogramming modules can be redeployed across cell types, offering new avenues for unlocking regenerative potential in otherwise non-regenerative tissues.

## Introduction

Injury- or disease-induced neuronal loss in the adult mammalian retina leads to irreversible visual impairment, owing to its lack of regenerative capacity. While no effective treatments currently exist to replenish lost retinal neurons, a promising approach is to reprogram endogenous Müller glia (MG) into functional neurons. MG in lower vertebrates mount a remarkable regenerative response following retinal injury by adopting embryonic retinal progenitor cell (RPC)-like properties that enable them to proliferate and regenerate lost retinal neurons.^1–5^ However, MG in mammals respond to retinal injury by undergoing reactive gliosis, thereby failing to re-enter the cell cycle or initiate neurogenesis.^5–9^ This raises the fundamental question of whether mammalian MG can be reprogrammed into an RPC-like state to serve as an endogenous source for neuronal regeneration.

Over the past decade, numerous studies have sought to reprogram mammalian MG by modulating genes that are differentially regulated in MG from regeneration-competent species. For instance, the activation of Wnt signaling,^10^ inhibition of the Hippo pathway,^11,12^ or manipulation of intrinsic cell cycle regulators^13^ promotes MG proliferation but fail to confer neurogenic competence. Alternatively, forced overexpression of the proneural transcription factor *Ascl1* combined with retinal injury and chromatin remodeling,^14–16^ simultaneous inhibition of Notch signaling and nuclear factor I transcription factors,^5,17^ or suppression of PROX1 transfer to MG^18^ promotes direct MG-to-neuron transdifferentiation at the cost of limited proliferation, raising concerns about the depletion of the endogenous MG pool. Although the sequential induction of MG proliferation followed by MG-derived neurogenesis has been achieved by combining existing approaches, their reliance on multiple genetic and pharmacological interventions presents significant barriers to clinical translation.^16,17,19,20^ Taken together, current mammalian MG reprogramming strategies inspired by inter-species comparisons have been informative but remain insufficient to couple proliferative and neurogenic competence, highlighting the need for a simpler approach that elicits a more complete regenerative response.

Since regenerative constraints are present throughout the mammalian central nervous system, a complementary approach may be to redeploy reprogramming paradigms identified in other mammalian neuronal lineages to MG. Hippocampal neural stem cells (NSCs), which share key functional and transcriptomic features with MG, exhibit a marked loss of proliferative and neurogenic competence with aging.^1,21,22^ We previously identified two age-associated regulators, *Plagl2* and *Dyrk1a*, whose simultaneous induction and inhibition (**i**nduction of ***<u>P</u>****lagl2* & **<u>a</u>**nti-***<u>D</u>****yrk1a*; iPaD), respectively, rejuvenate aged hippocampal NSCs *in vivo*, thereby restoring NSC self-renewal and neurogenic output, with accompanying cognitive improvement.^23^ Given the similarities between aged NSCs and MG, we asked whether iPaD could similarly rejuvenate adult mammalian MG toward a progenitor-like state capable of initiating a regenerative response.

Here, using an inducible transgenic mouse model, we demonstrate that *Plagl2* is sufficient to induce the rejuvenation of adult mouse MG into an RPC-like state with coupled proliferative and neurogenic competence *in vivo*. In the absence of retinal injury, *Plagl2*-activated MG migrated apically to the outer nuclear layer (ONL) and underwent multiple regulated rounds of cell division. Following N-methyl-D-aspartate (NMDA)-induced retinal injury, *Plagl2*-activated MG exhibited endogenous ASCL1 oscillations, a hallmark of multipotent embryonic NSCs, and initiated neurogenesis toward inner nuclear layer (INL) neuronal identities. Single-cell RNA sequencing (scRNA-seq) further confirmed that *Plagl2*-activated MG were reprogrammed into a late RPC-like state and progressively acquired neurogenic potential. At the functional level, multielectrode array recordings indicated that *Plagl2* induction partially restored light-evoked retinal ganglion cell (RGC) responses after NMDA injury. Finally, as a translational extension, we show that AAV-mediated *Plagl2* overexpression was sufficient to activate MG proliferation *in vivo*. Together, our work identifies *Plagl2* as a novel rejuvenating factor that unlocks the latent regenerative potential of adult mammalian MG, highlighting the broader concept that reprogramming modules may be redeployed across distinct cell types that otherwise lack intrinsic regenerative potential.

## Results

### *Plagl2* activates MG proliferation in the absence of retinal injury in adult mice

To assess the reprogramming potential of iPaD in adult mouse MG, we first examined whether its components, *Plagl2* and *Dyrk1a*, are endogenously regulated in MG following retinal injury. Analysis of a previously published scRNA-seq dataset of adult mouse retinas subjected to NMDA injury^5^ revealed that *Plagl2*, *Dyrk1a*, and the proneural transcription factor gene *Ascl1* were not differentially regulated in *Sox9*^+^ MG (Fig. S1A,B). *pHes5-iPaD-mCherry* transgenic mice, in which *Plagl2* cDNA and *Dyrk1a* short hairpin RNA are simultaneously expressed under the control of a doxycycline (Dox)-inducible Tet-ON system driven by an MG-specific *Hes5* promoter, were used to examine whether exogenous iPaD expression could reprogram adult mouse MG *in vivo* (Fig. 1A,B).^22,24,25^ Adult *pHes5-iPaD-mCherry* mice (∼2 months old) were treated with or without Dox for 1 week in combination with 5-Bromo-2′-deoxyuridine (BrdU) to label proliferating cells. Immunohistochemical analysis revealed that, in the absence of Dox, *pHes5-iPaD-mCherry* retinas showed minimal basal transgene activity, with no detectable PLAGL2 or mCherry signal in SOX9^+^ MG in the INL or SOX9^+^ astrocytes in the ganglion cell layer (GCL) (Fig. S1C–E).^26^ By contrast, Dox administration specifically induced PLAGL2 and mCherry expression in SOX9^+^ MG, with a subset incorporating BrdU and relocating apically into the ONL, recapitulating the interkinetic nuclear migration of embryonic RPCs and activated MG in regeneration-competent species (Fig. S1F–H, yellow arrowheads).^1,27^ In addition, consistent with prior work showing limited DYRK1A expression throughout the INL,^28^ DYRK1A was barely detectable in SOX2^+^ MG, even after NMDA injury (Fig. S1I–K). These data identify *Plagl2* as the dominant effector of the iPaD transgene in initiating MG proliferation; therefore, we focused subsequent analyses on *Plagl2* using the iPaD transgenic mice.

**Figure 1.**
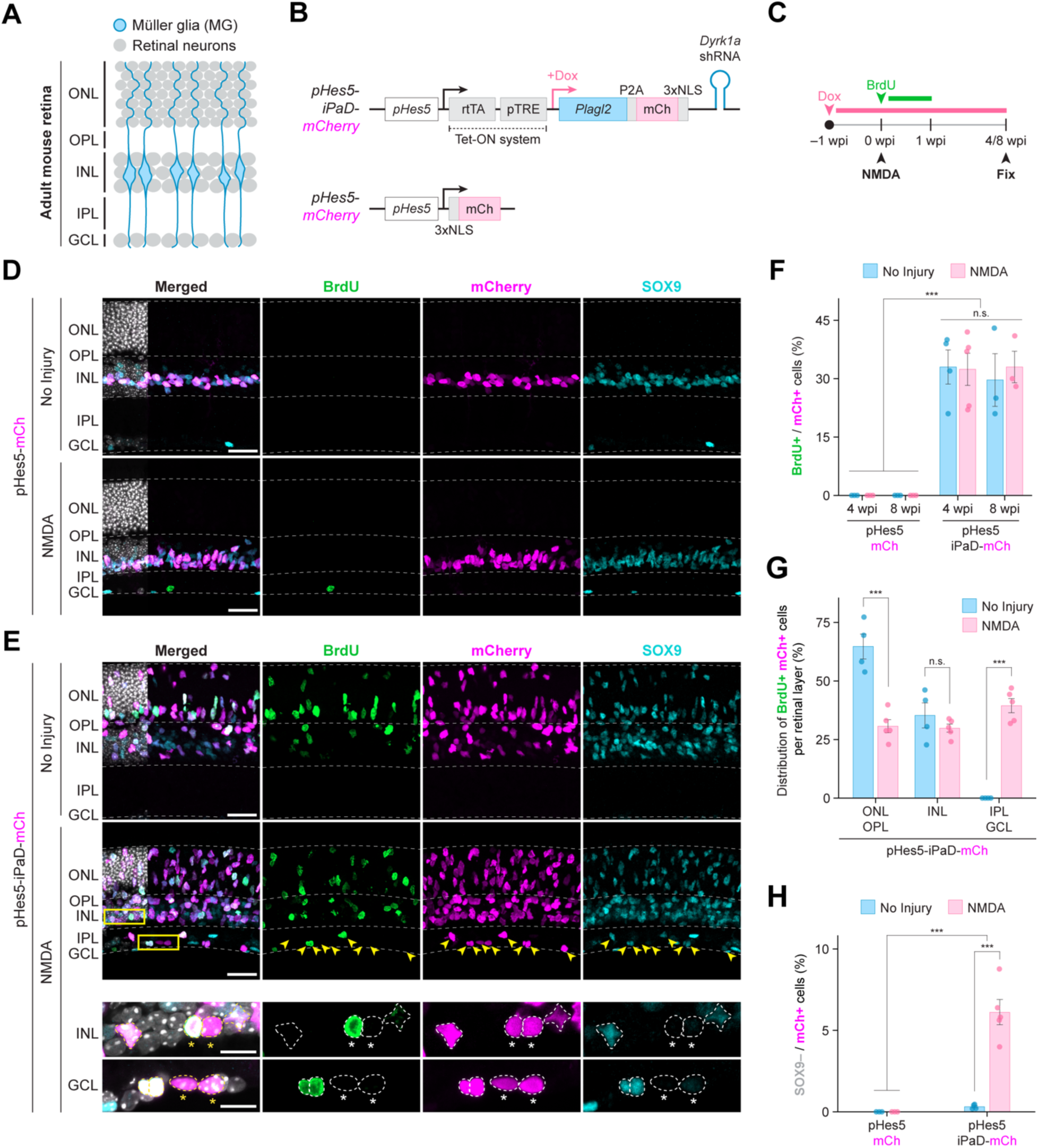
*Plagl2* activates MG proliferation without NMDA injury. A) Schematic of the adult mouse retina. GCL, ganglion cell layer; INL, inner nuclear layer; IPL, inner plexiform layer; ONL, outer nuclear layer; OPL, outer plexiform layer. B) Schematic of the *pHes5-iPaD-mCherry* and *pHes5-mCherry* transgene constructs. NLS, nuclear localization signal. C) Experimental paradigm to activate the iPaD transgene in adult mouse MG. D) Representative confocal images of *pHes5-mCherry* retinas at 4 wpi with or without NMDA injury. Sections were immunostained for BrdU (green), mCherry (magenta), and SOX9 (cyan). Scale bars, 25 µm. E) Representative confocal images of *pHes5-iPaD-mCherry* retinas at 4 wpi with or without NMDA injury. Sections were immunostained for BrdU (green), mCherry (magenta), and SOX9 (cyan). Yellow arrowheads indicate mCherry^+^ cells that migrated toward the GCL. Asterisks highlight examples of mCherry^+^/SOX9^−^ cells located within the INL and GCL. Boxed regions in the INL and GCL are enlarged in the bottom (zoomed) panels. Scale bars, 25 µm (main panels) and 10 µm (zoomed panels). F) Quantification of BrdU^+^/mCherry^+^ cells at 4 and 8 wpi. Data are presented as mean ± SEM. Significance was determined by three-way ANOVA followed by post hoc Tukey’s test. ***p < 0.001. n.s., not significant. *n* = 3–5 retinas per group. G) Distribution of mCherry^+^/BrdU^+^ cells across retinal layers in *pHes5-iPaD-mCherry* retinas at 4 wpi. Data are presented as mean ± SEM. Significance was determined by an unpaired *t-*test. ***p < 0.001. *n* = 4–5 retinas per group. H) Quantification of SOX9^−^/mCherry^+^ cells at 4 wpi. Data are presented as mean ± SEM. Significance was determined by two-way ANOVA followed by post hoc Tukey’s test. ***p < 0.001. *n* = 3–5 retinas per group.

We next investigated whether sustained *Plagl2* activation, in combination with NMDA injury, could further enhance MG proliferation. *pHes5-mCherry* mice, which express only mCherry under the *Hes5* promoter, were used as a control (Fig. 1B).^29^ To this end, *pHes5-iPaD-mCherry* and *pHes5-mCherry* mice were treated with Dox for 1 week prior to intravitreal NMDA injection, followed by an additional week of BrdU administration. Dox treatment was continued until retinal harvest at 4 and 8 weeks post-injury (wpi) (Figs. 1C–E and S2A). In control *pHes5-mCherry* retinas, mCherry expression was restricted to SOX9^+^ MG. Although NMDA injury led to a noticeable thinning of the inner plexiform layer, mCherry^+^/SOX9^+^ MG remained quiescent and confined to the INL at 4 wpi (Fig. 1D) and 8 wpi (Fig. S2A). Likewise, Dox-untreated *pHes5-iPaD-mCherry* retinas showed no evidence of MG proliferation or migration following NMDA injury (Fig. S2B). In contrast, Dox-treated *pHes5-iPaD-mCherry* retinas exhibited robust MG proliferation, with ∼30% of mCherry^+^ cells incorporating BrdU at both 4 and 8 wpi (Figs. 1E,F and S2A). While the overall extent of MG proliferation remained comparable between non-injured and injured retinas at both time points, NMDA injury significantly influenced the spatial distribution of MG-derived progeny. In non-injured retinas, 64.7 ± 5.3% of BrdU^+^/mCherry^+^ cells migrated apically toward the ONL, whereas following NMDA injury, 39.5 ± 3.0% of the progeny instead migrated basally toward the GCL, corresponding to the location of injury-induced neuronal loss (Fig. 1E, yellow arrowheads; Fig. 1G). Occasional BrdU^+^/mCherry^−^ cells were co-labeled with IBA1, identifying them as proliferating microglia (Fig. S2C,D). Notably, SOX9 expression was downregulated in a subset of mCherry⁺ cells within the INL and GCL (no injury, 0.30 ± 0.07%; NMDA injury, 6.0 ± 0.8%) and adopted a rounded, neuron-like morphology, suggesting the loss of glial identity and potential commitment to a neuronal fate (Fig. 1E, yellow boxes are enlarged in the bottom panels, and SOX9^−^/mCherry^+^ cells are indicated by asterisks; Fig. 1H). These results show that *Plagl2* drives injury-independent MG proliferation and apical nuclear migration, whereas NMDA injury redirects a subset of MG-derived progeny basally toward the GCL with the progressive attenuation of glial identity.

### *Plagl2*-driven MG proliferation proceeds through multiple rounds of regulated cell division

Given the robust proliferative response observed in *Plagl2*-activated MG, we examined the degree and dynamics of their re-entry into the cell cycle. To label successive rounds of cell division, Dox was administered to adult *pHes5-iPaD-mCherry* mice for 1 week, followed by the sequential administration of BrdU for 5 days and then 5′-Ethynyl-2′-deoxyuridine (EdU) for 5 days after a 2-day interval, enabling the discrimination of cells completing one or multiple rounds of cell division (Fig. 2A–C). Immunohistochemical analysis revealed that the majority of *Plagl2*-activated MG re-entered the cell cycle during the initial BrdU-labeling window in non-injured and NMDA-injured retinas, as only a few EdU^+^/mCherry^+^ cells (< 2%) lacked BrdU labeling (Fig. 2D). Notably, ∼20% of mCherry^+^ cells were double-labeled as BrdU^+^/EdU^+^ (no injury, 15.9 ± 3.0%; NMDA injury, 20.3 ± 1.0%), indicating S phase re-entry during the EdU-labeling window, whereas the single-labeled BrdU^+^/mCherry^+^ cells (no injury, 16.7 ± 3.3%; NMDA injury, 15.9 ± 1.9%) likely reflected cell cycle exit before the EdU-labeling window (Fig. 2D). Furthermore, the BrdU^+^/EdU^+^/mCherry^+^cells were predominantly located in the ONL in non-injured retinas, whereas 19.5 ± 7.0% migrated toward the GCL following NMDA injury (Fig. 2E). These data indicate that *Plagl2* generates transiently and persistently dividing MG populations, while NMDA injury primarily alters their spatial distribution rather than the extent of cell cycle re-entry.

**Figure 2.**
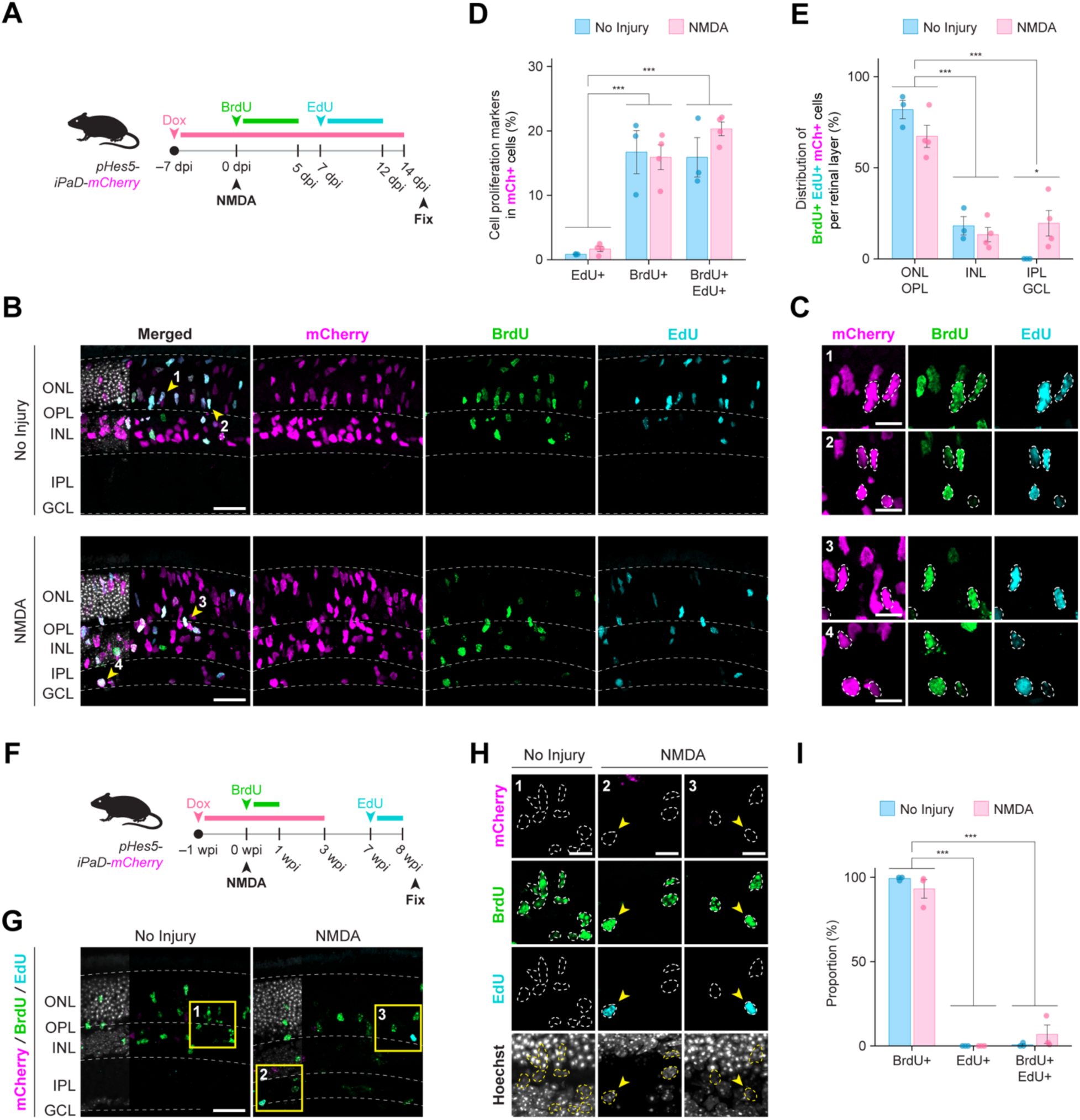
*Plagl2* induces multiple rounds of regulated cell divisions. A) Experimental paradigm to quantify MG cell cycle re-entry. B) Representative confocal images of *pHes5-iPaD-mCherry* retina at 14 dpi with or without NMDA injury. Sections were immunostained for BrdU (green), mCherry (magenta), and EdU (cyan). The areas indicated by arrowheads with numbers are enlarged in (C). Scale bar, 25 µm. C) Magnified images of BrdU^+^/EdU^+^/mCherry^+^ cells from (B), denoted by dotted outlines. Scale bar, 10 µm. D) Quantification of cell proliferation markers expressed in mCherry^+^ cells. Significance was determined by two-way ANOVA followed by post hoc Tukey’s test. ***p < 0.001. *n* = 3–4 retinas per group. E) Distribution of BrdU^+^/EdU^+^/mCherry^+^ cells across retinal layers. Significance was determined by two-way ANOVA followed by post hoc Tukey’s test. *p < 0.05, ***p < 0.001. *n* = 3–4 retinas per group. F) Experimental paradigm to assess oncogenic potential of reprogrammed MG. G) Representative confocal images from *pHes5-iPaD-mCherry* retina at 8 wpi with or without NMDA injury. Sections were immunostained for BrdU (green), mCherry (magenta), and EdU (cyan). Boxed regions are enlarged in (H). Scale bar, 25 µm. H) Magnified images of BrdU^+^/EdU^+^ cells from (G). Scale bar, 10 µm. I) Quantification of BrdU and EdU uptake. Significance was determined by two-way ANOVA followed by post hoc Tukey’s test. ***p < 0.001. *n* = 3 retinas per group.

We next investigated whether *Plagl2*-activated MG remained under regulatory control or were susceptible to tumorigenic expansion. To assess the persistence of proliferative activity after *Plagl2* withdrawal, Dox was administered to adult *pHes5-iPaD-mCherry* mice for 1 week, followed by NMDA injury and 1 week of BrdU labeling. Dox treatment was discontinued at 3 wpi, and EdU was administered during the final week before retinal harvest at 8 wpi (Fig. 2F). Immunohistochemical analysis confirmed the loss of mCherry expression by 8 wpi, indicating successful transgene shutdown (Fig. 2G,H). Despite this, BrdU^+^ cells persisted in non-injured and NMDA-injured retinas, with a spatial distribution similar to that observed during active transgene expression (Fig. 1E). Notably, only a minor fraction of BrdU^+^ cells incorporated EdU (no injury, 0.6 ± 0.6%; NMDA injury, 6.9 ± 5.5%), and none of them exhibited tumor-like cell clusters (Fig. 2G,H, boxes 2–3; Fig. 2I). These findings demonstrate that *Plagl2*-activated MG undergo multiple rounds of cell division in a self-limiting manner, with minimal evidence of tumorigenic expansion following the cessation of transgene expression.

### *Plagl2* induces ultradian ASCL1 oscillations in MG following NMDA injury

In the aged mouse hippocampus, iPaD rejuvenates quiescent NSCs by relieving age-associated chromatin constraints at the *Ascl1* locus to induce oscillatory ASCL1 expression with a periodicity of 2–3 h, a dynamic transcriptional hallmark observed in multipotent embryonic NSCs.^23,30^ In parallel, regeneration-competent zebrafish MG rapidly induce *ascl1a* expression following injury to initiate neurogenesis, whereas mammalian MG fail to mount comparable *Ascl1* upregulation and instead enter a reactive gliotic program that returns toward quiescence by 72 h post injury (hpi).^3–5,12^ To gain a mechanistic insight into how *Plagl2* drives MG reprogramming, we asked whether *Plagl2* induces ASCL1 expression in MG during this early injury-response phase at 24, 48, and 72 hpi (Fig. 3A). Immunohistochemical analysis showed that ASCL1 expression remained undetectable in control *pHes5-mCherry* retinas across all time points, consistent with the scRNA-seq data (Figs. 3B,C, S1B, and S3A,B). By contrast, *Plagl2*-activated MG showed the progressive induction of ASCL1 following NMDA injury, reaching 22.3 ± 1.7% at 72 hpi (Fig. 3C).

**Figure 3.**
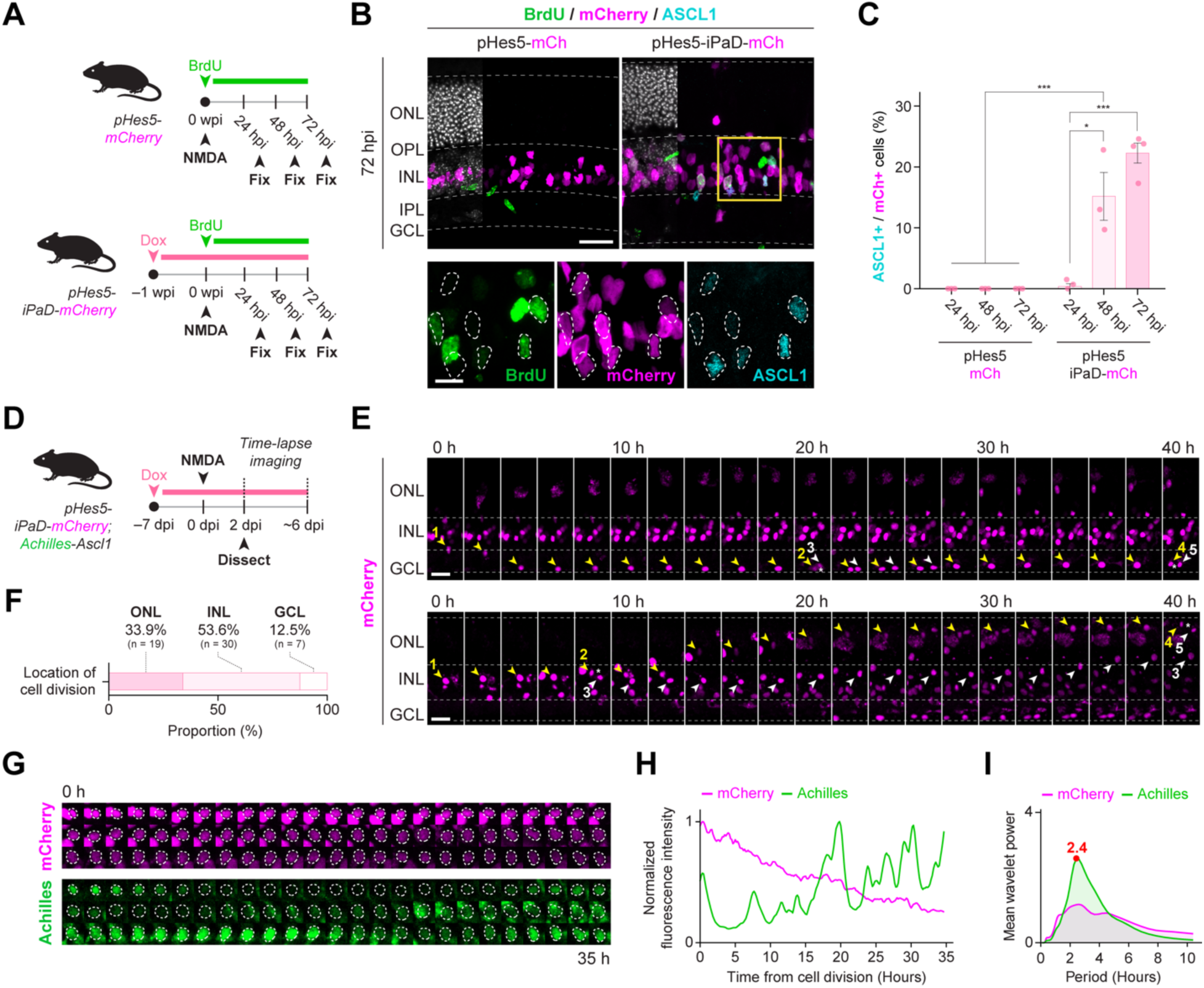
*Plagl2* induces ultradian ASCL1 oscillations in MG following NMDA injury. A) Experimental paradigm for time-course analysis following NMDA injury. B) Representative confocal images of *pHes5-mCherry* and *pHes5-iPaD-mCherry* retinas at 72 hpi. Sections were immunostained for BrdU (green), mCherry (magenta), and ASCL1 (cyan). Examples of ASCL1^+^/mCherry^+^ cells denoted by dotted outlines are shown below. Scale bars, 25 µm (main panels) and 10 µm (zoomed panels). C) Quantification of ASCL1^+^/mCherry^+^ cells from (B). Data are presented as mean ± SEM. Significance was determined by two-way ANOVA followed by post hoc Tukey’s test. ***p < 0.001. *n* = 3–4 retinas per group. D) Experimental paradigm for *ex vivo* retinal slice culture and time-lapse imaging. E) Representative confocal time-lapse montages showing *Plagl2*-activated MG dividing in the INL and migrating toward the ONL/GCL. Arrowheads mark mother, daughter, and granddaughter cells. White asterisks denote cell division. Scale bar, 25 µm. F) Quantification of cell division occurring across retinal layers from (E). *n* = 3 retinas. G) Representative single-cell track from time-lapse imaging of *Plagl2*-activated MG showing *Achilles-Ascl1* and *iPaD-mCherry* expression. H) Quantification of fluorescence intensity from the representative single-cell track in (G). I) Mean wavelet power spectra for Achilles and mCherry fluorescence. *n* = 25 tracks from 3 retinas.

To examine whether *Plagl2*-activated MG show ASCL1 oscillations, *pHes5-iPaD-mCherry* mice were crossed with *Achilles-Ascl1* knock-in reporter mice, in which the fluorescent protein Achilles is fused in-frame to the 5′ end of the endogenous *Ascl1* locus (Fig. S3C–E). While *Ascl1*-null mice die shortly after birth, homozygous *Achilles-Ascl1* mice were fertile without severe phenotypic defects, suggesting that *Achilles-Ascl1* protein was functional. Immunohistochemical analysis of embryonic *pHes5-iPaD-mCherry*;*Achilles-Ascl1* retinas, a stage at which ASCL1 is endogenously expressed,^31,32^ confirmed that Achilles fluorescence overlapped with endogenous ASCL1 protein, validating the proper integration and expression of the Achilles knock-in reporter (Fig. S3F,G). To validate further its ability to capture ASCL1 oscillations, time-lapse imaging of primary RPCs derived from embryonic *pHes5-iPaD-mCherry;Achilles-Ascl1* retinas was performed and the periodicity of Achilles fluorescence was quantified using wavelet transform analysis. This analysis revealed oscillatory ASCL1 expression in RPCs with an average periodicity of ∼2 h, consistent with that observed in embryonic NSCs (Fig. S3H,I).^30^

*Ex vivo* time-lapse imaging of adult *pHes5-iPaD-mCherry;Achilles-Ascl1* retinal explants following Dox treatment and NMDA injury to track ASCL1 expression dynamics in *Plagl2*-activated MG (Fig. 3D) revealed the nuclear migration of mCherry^+^ cells from the INL to either the ONL or GCL, with a subset undergoing two rounds of cell division (Fig. 3E arrowheads, F; Movie S1). Strikingly, in a subset of mCherry^+^ cells that divided in the INL, ASCL1 oscillated with an average periodicity of 2.4 h, recapitulating the oscillations observed in RPCs and embryonic NSCs (Fig. 3G–I; Movie S2). By contrast, mCherry expression remained stable or declined gradually over time, showing no evidence of periodicity, consistent with the downregulation of *Hes5* activity during neurogenic reprogramming (Figs. 3G,H and S3J). Taken together, these findings indicate that *Plagl2* induces ultradian ASCL1 oscillations in MG during the early post-injury phase, thereby supporting their proliferation while priming them for the acquisition of neurogenic competence.

### *Plagl2*-activated MG initiate neurogenesis toward INL neurons upon NMDA injury

We next asked whether *Plagl2*-induced neurogenic competence gives rise to defined mature retinal neuronal identities. Using the same induction and injury paradigm described in Fig. 1C, immunohistochemistry for OTX2 and NEUN, which label bipolar/photoreceptor (PR) and amacrine/RGC fates, respectively,^5,17^ was performed. In control *pHes5-mCherry* retinas with or without injury, mCherry^+^ MG were mutually exclusive with OTX2^+^ and NEUN^+^ neurons (Fig. S4A,B), indicating that MG do not obtain neurogenic competence even after NMDA injury. By contrast, although in non-injured *pHes5-iPaD-mCherry* retinas, <1% of BrdU^+^/mCherry^+^ cells co-expressed neuronal markers, NMDA injury significantly increased the fraction co-expressing OTX2 (5.0 ± 1.2%) and NEUN (5.8 ± 1.2%) (Fig. 4A–C). Furthermore, the proportion of BrdU^+^/mCherry^+^/OTX2^+^ cells increased significantly from 4 to 8 wpi (11.0 ± 2.4%), suggesting that prolonged *Plagl2* expression enhances MG-derived neurogenesis toward a bipolar fate (Fig. 4C). Although a subset of BrdU^+^/mCherry^+^ cells co-expressed OTX2 in the ONL, they lacked PR-like nuclear morphology and did not express the PR marker recoverin (Fig. 4B, box 1; Fig. S4D,E). The proportion of BrdU^−^/mCherry^+^ cells co-expressing neuronal markers, which may represent MG that either proliferated outside the BrdU-labeling window or underwent direct transdifferentiation, was also quantified. In *pHes5-iPaD-mCherry* retinas subjected to NMDA injury, this population declined from 4 wpi (OTX2 = 4.2 ± 0.8%; NEUN = 5.9 ± 0.6%) to 8 wpi (OTX2 = 2.2 ± 1.0%; NEUN = 3.5 ± 0.9%), likely reflecting the loss of *Hes5*-driven mCherry expression as reprogrammed MG progressively acquired mature neuronal identities (Fig. S4C). Consistently, mCherry^+^ cells that co-expressed neuronal markers exhibited reduced mCherry intensity compared with those that did not, suggesting that MG-derived neuronal output may be underestimated by *Hes5* promoter-based lineage labeling (Fig. S4F). Nonetheless, we additionally confirmed that in *pHes5-iPaD-mCherry* retinas following NMDA injury, ∼10% of BrdU^+^/mCherry^+^ cells in the INL and GCL co-expressed the amacrine/RGC marker HUC/D (4 wpi, 8.3 ± 0.5%; 8 wpi, 7.5 ± 0.8%), whereas none co-expressed the mature RGC marker BRN3A in *pHes5-iPaD-mCherry* (Fig. 4D–F) and *pHes5-mCherry* retinas (Fig. S4G,H). Overall, these results suggest that *Plagl2* alone induces rare neurogenesis toward an OTX2^+^ bipolar fate, whereas NMDA injury markedly enhances neurogenic competence toward INL neuronal identities, including OTX2^+^ bipolar and NEUN^+^/HUC/D^+^ amacrine fates.

**Figure 4.**
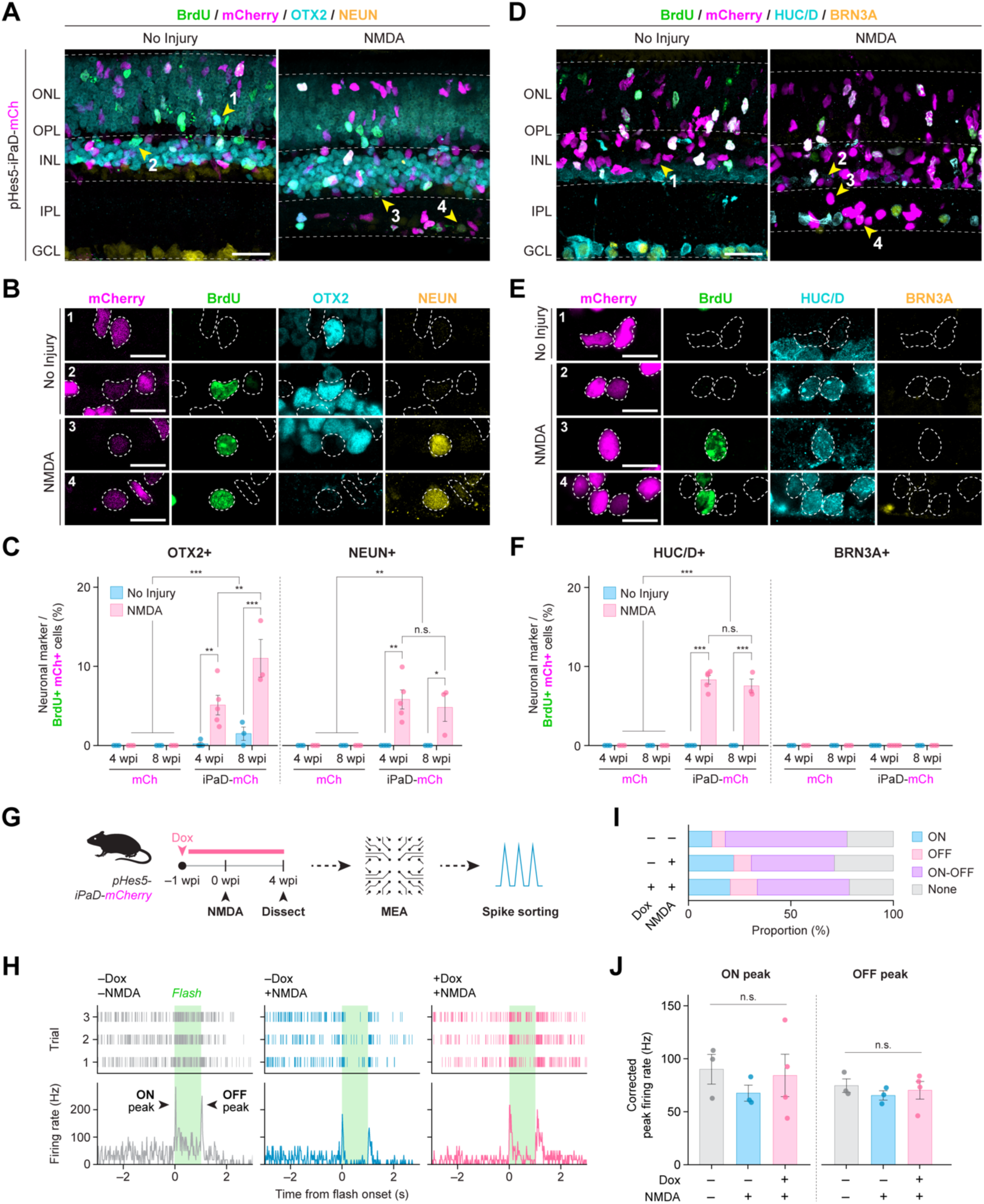
*Plagl2*-activated MG initiate neurogenesis toward INL neurons upon NMDA injury. A) Representative confocal images from *pHes5-iPaD-mCherry* retinas at 4 wpi with or without NMDA injury. Sections were immunostained for BrdU (green), OTX2 (cyan), and NEUN (yellow). mCherry signal (magenta) was detected by native fluorescence. Scale bars = 25 µm. B) Magnified images of mCherry^+^ cells from (A). Scale bar, 10 µm. C) Quantification of BrdU^+^/mCherry^+^ cells that co-express OTX2 or NEUN. Data are presented as mean ± SEM. Significance was determined by three-way ANOVA followed by post hoc Tukey’s test. *p < 0.05, **p < 0.01, ***p < 0.001. *n* = 3–5 retinas per group. D) Representative confocal images of control *pHes5-iPaD-mCherry* retinas at 4 wpi with or without NMDA injury. Sections were immunostained for BrdU (green), mCherry (magenta), HUC/D (cyan), and BRN3A (yellow). Scale bars, 25 µm. E) Magnified images of mCherry^+^ cells from (D). Scale bar, 10 µm. F) Quantification of BrdU^+^/mCherry^+^ cells that co-expressing HUC/D or BRN3A. Data are presented as mean ± SEM. Significance was determined by three-way ANOVA followed by post hoc Tukey’s test. ***p < 0.001. *n* = 3–5 retinas per group. G) Experimental paradigm for multielectrode array (MEA) recording and analysis H) Representative ON-OFF light responses from each experimental condition. For each unit, spike rasters (top) and peri-stimulus time histograms (bottom) are shown. The light stimulus period (0–1 s) is indicated by green shading. I) Proportion of RGC response classes across experimental conditions after spike sorting. J) Mouse-level comparison of corrected ON and OFF peak firing rates across experimental conditions. The ON peak panel was calculated from units reclassified as ON or ON-OFF, whereas the OFF peak panel was calculated from units reclassified as OFF or ON-OFF. *n* = 3–4 mice per group. Statistical comparisons were performed using the Kruskal–Wallis test.

To assess further whether MG-derived neurons are functional and contribute to the recovery of visual function following injury, multielectrode array recordings were used to measure RGC output from whole-mount retinas of *pHes5-iPaD-mCherry* mice with or without NMDA injury (Fig. 4G). For each unit, spike rasters and peri-stimulus time histograms were generated from RGC responses evoked by a 1-s flash stimulus under dark-adapted conditions (Fig. 4H). The responses were classified after spike sorting as “ON,” “OFF,” “ON-OFF,” or “none,” and the corrected peak firing rates of the ON and OFF components were quantified as described previously (Fig. 4I).^33,34^ Although the overall distribution of RGC response classes was comparable across experimental conditions, the corrected ON and OFF peak firing rates showed modest differences (Figs. 4J and S4I). In control *pHes5-iPaD-mCherry* retinas (–Dox; –NMDA), robust ON and OFF responses were observed. NMDA injury (–Dox; +NMDA) significantly reduced the firing rates of both response types, consistent with impaired retinal function following inner retinal damage. Notably, in *Plagl2*-induced retinas with NMDA injury (+Dox; +NMDA), the ON and OFF responses partially shifted toward control levels, although there were no significant differences between the experimental conditions for either peak response. These findings nevertheless suggest a trend toward improved light-evoked RGC output following *Plagl2* induction in injured retinas, consistent with a possible functional contribution from MG-derived neurons.

### scRNA-seq reveals progressive neurogenic reprogramming of MG following *Plagl2* induction

To define the transcriptomic profile of MG following *Plagl2* induction, scRNA-seq was performed on fluorescence-activated cell sorting-purified mCherry^+^;DAPI^−^ cells from NMDA-injured control *pHes5-mCherry* retinas and *pHes5-iPaD-mCherry* retinas with or without NMDA injury at 4 wpi (Fig. 5A). After quality control, 15,410 (*pHes5-mCherry*; +NMDA), 10,109 (*pHes5-iPaD-mCherry*; no injury), and 11,880 (*pHes5-iPaD-mCherry*; +NMDA) high-quality cells were integrated and projected onto a shared uniform manifold approximation and projection (UMAP) embedding (Figs. 5B and S5A). *Plagl2* induction resulted in the reduced expression of the canonical MG marker *Glul*, whereas endogenous *Plagl2* and *Dyrk1a* expression remained low overall, with modest increases in the fraction of positive cells (Fig. S5B). By contrast, the neurogenic regulators *Ascl1* and *Neurog2* were upregulated and detected in a larger fraction of cells, supporting transcriptional remodeling following *Plagl2* induction (Fig. S5B). Unsupervised clustering resolved 14 clusters, which were annotated into broader cell classes using established marker genes (Figs. 5C and S5C; Table S1). The MG cluster was defined by the high expression of canonical MG marker genes (*Glul*, *Aqp4*, *Sox9*, *Rlbp1*, and *Slc1a3*) and was predominantly composed of cells from control *pHes5-mCherry* retinas (Fig. 5D,E). Notably, *Plagl2* induction generated distinct reprogrammed populations: a reprogrammed MG cluster with reduced MG marker expression and increased progenitor-associated gene expression (*Hes5*, *Ccnd1*, *Ccnd2*, and *Nes*), a dividing MG cluster enriched for S/G2M-phase genes (*Cenpf*, *Mki67*, and *Top2a*), and a neurogenic MG cluster expressing pan-neurogenic regulators (*Ascl1*, *Neurog2*, and *Insm1*) as well as neuronal markers (*Rbfox3/NeuN*, *Elavl3/HuC*, and *Tubb3*) (Figs. 5B–E and S5D). We also identified neuron-like clusters, including a bipolar-like cluster expressing mature bipolar markers (*Otx2*, *Lhx4*, and *Cabp5*) and two PR-like clusters. PR-like-1 was enriched for mature PR markers (*Rho*, *Rcvrn*, *Gnat1*, and *Gnat2*), whereas PR-like-2 was enriched for early PR markers (*Crx*, *NeuroD1*, and *Prdm1*) (Fig. S5E). Although PR-like-2 was transcriptionally distinct from MG and could be interpreted as *bona fide* MG-to-PR conversion, both PR-like clusters were also present in control *pHes5-mCherry* retinas and resembled the PR-like signatures reported previously as technical artifacts (Fig. S5F,G).^13,15,35^ By contrast, the neurogenic MG and bipolar-like populations were absent from control *pHes5-mCherry* retinas and became enriched in *pHes5-iPaD-mCherry* retinas with NMDA injury, consistent with immunohistochemical results showing the enhanced acquisition of neurogenic competence toward INL identities following NMDA injury (Figs. 5D,E and S5F). These observations suggest that *Plagl2* induction drives broad transcriptional reprogramming in MG and that the neurogenic MG and bipolar-like populations represent *bona fide* MG-derived neurogenic conversion trajectories.

**Figure 5.**
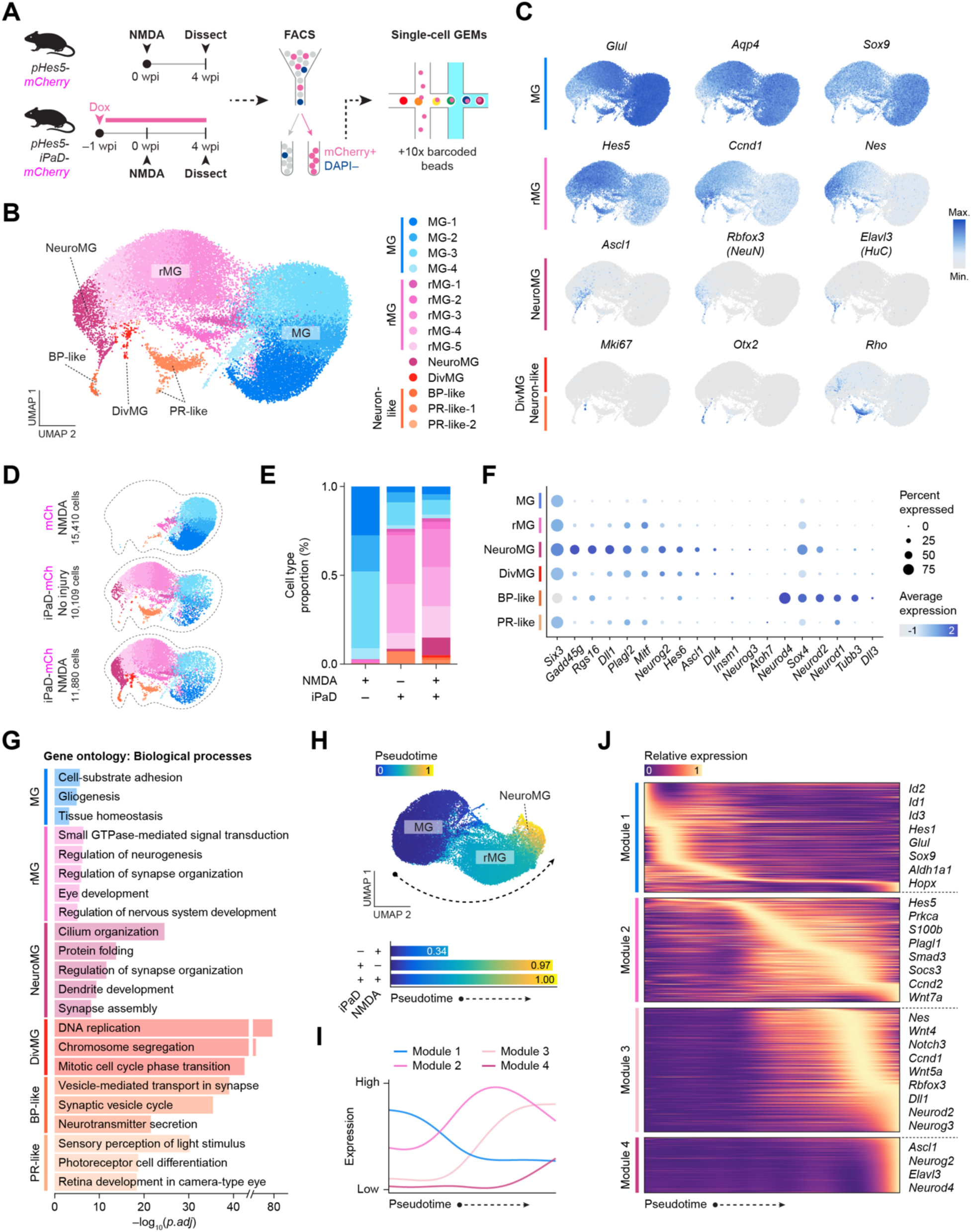
scRNA-seq reveals progressive neurogenic reprogramming of MG following *Plagl2* induction. A) Experimental paradigm for scRNA-seq analysis. B) Integrated UMAP of fluorescence-activated cell sorting (FACS)-purified *pHes5-mCherry* and *pHes5-iPaD-mCherry* retinas with cluster annotations. GEMS, gel beads-in-emulsion. C) Expression of representative marker genes for MG, reprogrammed MG (rMG), neurogenic MG (NeuroMG), dividing MG (DivMG), and neuron-like clusters. D) Integrated UMAP split by sample identity. E) Quantification of annotated cell type proportion<u>s</u> per sample identity. F) Dot plot of representative marker genes involved in retinal development and neurogenesis. G) Gene ontology enrichment for annotated clusters. H) Pseudotime analysis of the MG–rMG–NeuroMG trajectory. I) Mean expression of each gene module across pseudotime. J) Heatmap of representative module genes ordered by pseudotime.

Given the upregulation of progenitor-associated genes in the reprogrammed MG and neurogenic MG clusters, we asked whether *Plagl2* rejuvenates MG by reactivating a developmental RPC-like transcriptional program. To address this, transfer anchors were identified against a previously annotated mouse retinal developmental scRNA-seq reference and our query cells were projected onto the reference UMAP embedding (Fig. S5H–J). Cells from control *pHes5-mCherry* retinas mapped predominantly to the reference MG compartment. By contrast, cells from non-injured *pHes5-iPaD-mCherry* retinas showed a redistribution toward late RPCs and amacrine identities, whereas NMDA injury further increased cell assignment to late RPCs and bipolar identities with high prediction scores, accompanied by expression of late RPC/neurogenic-associated genes (*Gadd45g, Rgs16, Dll1, Sox4,* and *Hes6*) (Figs. 5F and S5H–J).^32^ Together, these results demonstrate that *Plagl2* rejuvenates adult MG toward a late RPC-like state with bipolar-and amacrine-biased neurogenic potential.

To interpret these diverse cell states functionally, gene ontology (GO) enrichment analysis was performed across the annotated clusters (Fig. 5G). The MG cluster was enriched for terms involved in glial homeostatic processes including cell-substrate adhesion, gliogenesis, and tissue homeostasis. In contrast, the reprogrammed MG cluster was enriched for signaling and developmental programs, including small GTPase-mediated signal transduction, eye development, and neurogenesis. The neurogenic MG cluster was further enriched for neuronal maturation and synaptic development, with cilium organization emerging as an unexpected category. The dividing MG, bipolar-like, and PR-like clusters were enriched for their corresponding functional terms, including cell cycle progression, neuronal transmission, and visual function, respectively. To confirm whether these diverse state-specific programs were under direct *Plagl2* transcriptional control, the genes upregulated in *Plagl2*-activated MG were compared with PLAGL2-chromatin immunoprecipitation-sequencing (ChIP-seq) targets from NSCs (Fig. S5K,L; Table S2).^23^ This identified an overlap of 897 genes that were targeted by PLAGL2 in NSCs and upregulated in *Plagl2*-activated MG regardless of injury status, and enriched for genes involved in neurogenesis including *Ascl1* and *Neurog2*, synapse formation/organization, and canonical Wnt signaling, which has been shown to be activated by *Plagl2*.^23,36,37^ In addition, 1,483 genes were upregulated in *Plagl2*-activated MG under both injury conditions yet absent from the ChIP-seq target list, which included cilium-associated and eye development terms. These data position *Plagl2* as a conserved upstream driver of neurogenic reprogramming, while revealing an additional MG-specific, context-dependent transcriptional module.

Having defined these MG-derived transcriptional states, we asked whether *Plagl2* induction drives a continuous reprogramming trajectory. Pseudotime analysis of the MG, reprogrammed MG, and neurogenic MG clusters supported a model in which the MG cluster first underwent a relatively discrete shift toward reprogrammed MG, followed by a continuous progression from reprogrammed MG to neurogenic MG (Fig. 5H). Coordinated gene programs were then identified by clustering trajectory-associated genes with shared dynamics along pseudotime, which resolved four major gene modules (Fig. 5I,J). Module 1 was initially highly expressed and progressively declined along pseudotime, comprising glia-associated genes including *Hes1* and *Id* family genes. Modules 2 and 3 rose sequentially after the initial glial phase and captured intermediate progenitor-activation and neurogenic transition programs including *Ccnd1* and *Dll1*. Module 4 emerged terminally and marked neurogenic commitment, and was enriched for core neurogenic regulators such as *Ascl1* and *Neurog2*. Together, these data identify *Plagl2* as a driver of continuous MG reprogramming through progenitor-like intermediate states toward neurogenic commitment, with NMDA injury reinforcing a late RPC-like state biased toward INL identities.

### Adeno-associated virus (AAV)-mediated *Plagl2* overexpression induces MG proliferation

Having demonstrated that *Plagl2* activates MG with coupled proliferative and neurogenic competence in the iPaD transgenic mouse line, and since *Dyrk1a* expression is minimal in MG, we next sought to assess the translational potential of *Plagl2* using an AAV-based delivery system. AAV constructs were generated that expressed either mCherry alone or *Plagl2-P2A-mCherry*, both lacking a nuclear localization signal on mCherry to allow the visualization of full cell morphology, under the control of a shortened 765-bp *Hes5* promoter, which was used due to a packaging limitation (Fig. 6A). The AAV2.7m8 capsid was used to target MG efficiently, and two tyrosine-to-phenylalanine substitutions (Y444F and Y730F) were introduced to enhance transduction.^38–40^ AAV was injected intravitreally into adult wild-type mice, followed by NMDA injection 1 week later, and retinas were harvested at 4 wpi (Fig. 6B). BrdU was administered for 2 weeks, beginning at the time of AAV injection, to maximize the labeling of MG-derived progeny.

**Figure 6.**
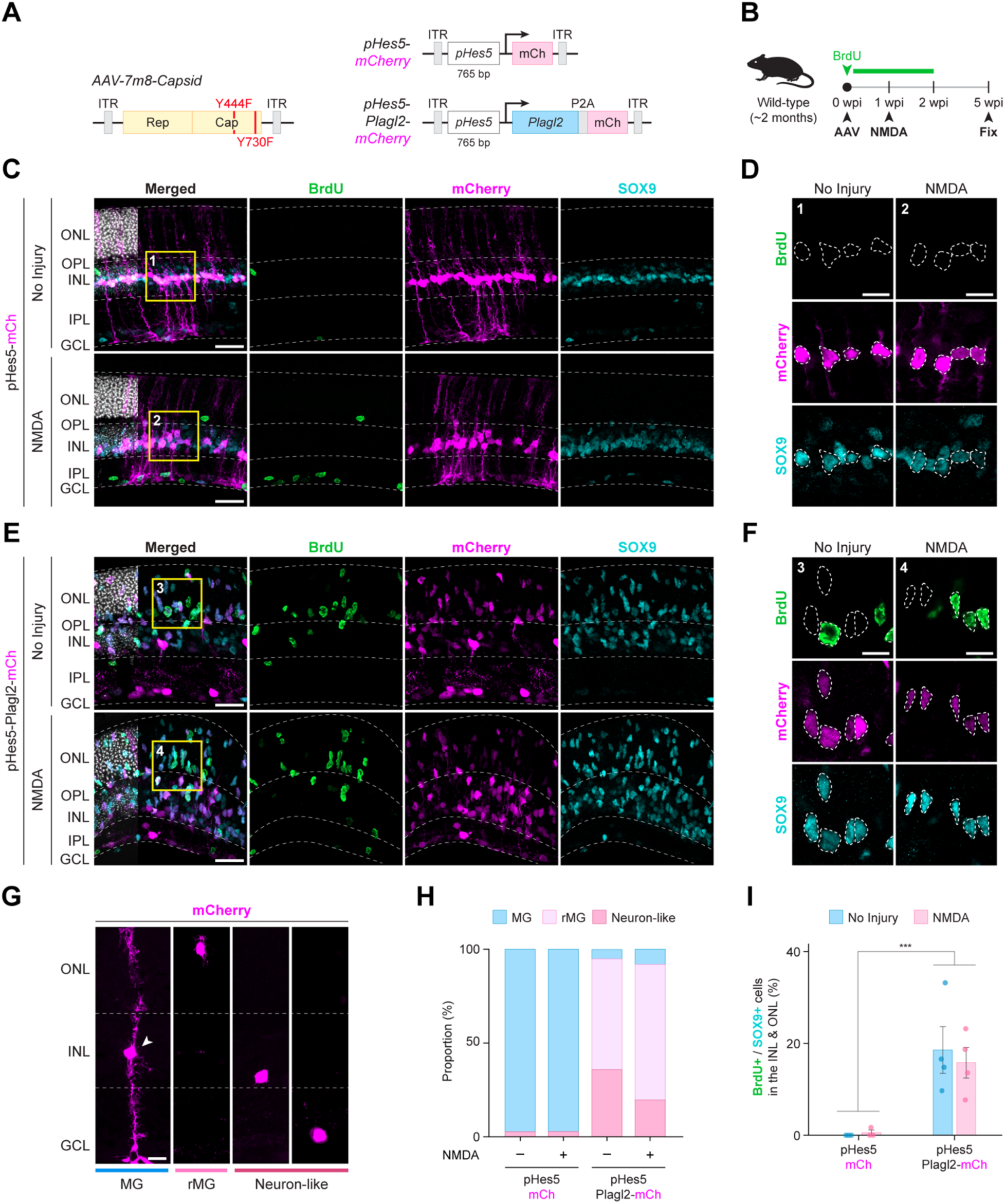
AAV-mediated *Plagl2* overexpression induces MG proliferation. A) Schematic of the AAV2.7m8 capsid, *AAV-pHes5-mCherry*, and *AAV-pHes5-Plagl2-mCherry* constructs. B) Experimental paradigm for AAV delivery. C) Representative confocal images of retinas infected with *AAV-pHes5-mCherry* with or without NMDA injury. Sections were immunostained for BrdU (green), mCherry (magenta), and SOX9 (cyan). Scale bars, 25 µm. D) Magnified images of mCherry^+^ cells from (C). Scale bar, 10 µm. E) Representative confocal images of retinas infected with *AAV-pHes5-Plagl2-mCherry* with or without NMDA injury. Sections were immunostained for BrdU (green), mCherry (magenta), and SOX9 (cyan). Scale bars, 25 µm. F) Magnified images of mCherry^+^ cells from (E). Scale bar, 10 µm. G) Representative images of mCherry^+^ cells with distinct cell morphologies. Scale bar, 10 µm. H) Quantification of the relative proportion of MG, reprogrammed MG (rMG) and neuron-like mCherry^+^ cells from retinas injected with *AAV-pHes5-mCherry* and *AAV-pHes5-Plagl2-mCherry* with or without NMDA injury. *n* = 3–4 retinas per group. I) Quantification of BrdU^+^/SOX9^+^ cells in the INL and ONL. Data are presented as mean ± SEM. Significance was determined by two-way ANOVA followed by post hoc Tukey’s test. ***p < 0.001. *n* = 3–4 retinas per group.

To verify transgene expression, PLAGL2 immunoreactivity was first examined in control (*AAV-mCherry*) and *Plagl2*-overexpressing (*AAV-Plagl2*) retinas. As expected, mCherry^+^ cells in control *AAV-mCherry* retinas remained confined to the INL, whereas many mCherry^+^ cells in *AAV-Plagl2* retinas migrated apically to the ONL under the injury and no injury conditions (Figs. 6C,E and S6A,B), resembling the transgenic phenotype. Correspondingly, PLAGL2 expression was absent from control *AAV-mCherry* retinas, but was readily detected in mCherry^+^ cells within the INL and ONL of *AAV-Plagl2* retinas under both injury conditions (Fig. S6A,B). We next examined the identity and morphology of mCherry^+^ cells. In control *AAV-mCherry* retinas, mCherry expression remained confined to SOX9^+^ MG in the INL, with elongated glial processes spanning the retinal layers (Fig. 6C,D). By contrast, *AAV-Plagl2* retinas contained many mCherry^+^/SOX9^+^ cells that retracted their glial processes and adopted a progenitor-like morphology (Fig. 6E,F).^17^ The fraction of mCherry^+^/SOX9^+^ cells was similar between *AAV-mCherry* and *AAV-Plagl2* retinas, indicating comparable MG transduction (Fig. S6C). To quantify the associated morphological shift, mCherry^+^/SOX9^+^ cells with glial processes were classified as MG, mCherry^+^/SOX9^+^ cells with retracted glial processes were classified as reprogrammed MG, and mCherry^+^/SOX9^−^ cells were classified as neuron-like (Fig. 6G,H). This revealed that control *AAV-mCherry* retinas were composed almost entirely of MG (no injury, 97.3 ± 0.1%; NMDA injury, 97.2 ± 0.1%), whereas mCherry^+^ cells in *AAV-Plagl2* retinas were composed predominantly of reprogrammed MG (no injury, 59.0 ± 7.0%; NMDA injury, 72.3 ± 3.1%) and neuron-like (no injury, 35.8 ± 8.3%; NMDA injury, 19.6 ± 3.9%) (Fig. 6H). A small number of neuron-like cells were also present in control *AAV-mCherry* retinas, indicating that AAV leakage into endogenous neurons occurred in subsets of cells, as described previously.^41^ However, neuron-like cells were dramatically increased in number in *AAV-Plagl2* retinas, suggesting that MG-derived neurogenesis was activated by *Plagl2*. We then asked whether this *Plagl2*-induced morphological transition was accompanied by

MG proliferation. Consistent with the transgenic model, BrdU incorporation was readily detected in many SOX9^+^ cells in *AAV-Plagl2* retinas within the INL and ONL, regardless of injury status (no injury, 18.6 ± 5.1%; NMDA injury, 15.8 ± 3.3%), but not in *AAV-mCherry* retinas (Fig. 6I). In parallel, the number of SOX9^+^ cells in the INL and ONL increased by ∼1.5-fold in *AAV-Plagl2* retinas compared to control *AAV-mCherry* retinas. Furthermore, many SOX9^+^ cells that migrated apically to the ONL showed faint or no mCherry expression, suggesting the dilution of the AAV signal due to cell division (Fig. 6E,F). Lastly, because mCherry fluorescence became faint in proliferating progeny, we assessed whether AAV-mediated *Plagl2* overexpression was sufficient to confer neurogenic competence by examining BrdU^+^ cells in *AAV-Plagl2* retinas for the co-expression of neuronal markers. Although rare, BrdU^+^/OTX2^+^ cells were detected in non-injured and injured *AAV-Plagl2* retinas, whereas there were no BrdU^+^/NEUN^+^ cells were detected (Fig. S6E,F). Together, these results indicate that AAV-mediated *Plagl2* overexpression is sufficient to drive robust MG proliferation with limited neurogenic competence, including rare acquisition of OTX2^+^ bipolar-like features, regardless of injury status.

## Discussion

Restoring lost neuronal function in the adult retina remains a central challenge in the field of regenerative neuroscience. Here, we show that *Plagl2* unlocks the latent regenerative potential of adult mouse MG by coupling injury-independent proliferation with injury-dependent neurogenesis. This coupled response distinguishes *Plagl2*-driven MG reprogramming from prior strategies, such as injury-dependent neurogenic paradigms with limited proliferation^5,14,35^ and injury-independent proliferative paradigms that yield little or no neurogenesis even following injury.^10–13^

The ability of *Plagl2* to coordinate these distinct regenerative phases likely stems from its conserved role as a developmental regulator of growth. *Plagl2* belongs to the PLAG family of zinc-finger transcription factors (*Plagl1/Zac1*, *Plag1*, and *Plagl2*) that regulate growth control and tumorigenesis across diverse cellular contexts. *Plagl1/Zac1* is associated with tumor-suppressive programs, whereas *Plag1* and *Plagl2* contribute to telencephalic progenitor expansion^42^ and proto-oncogenic activity.^37,43^ In the context of retinal development, *Plagl1/Zac1* has been implicated as a growth-suppressive regulator that promotes the cell cycle exit of RPCs to facilitate MG differentiation.^44^ By contrast, we found that *Plagl2* expression in MG did not induce tumorigenesis *in vivo*, but instead drove regulated rounds of cell cycle re-entry with limited neurogenesis, and this neurogenic output was markedly enhanced after NMDA injury. Similarly, in aged hippocampal NSCs, *Plagl2* alone promotes robust NSC proliferation, but yields limited neurogenesis, whereas *Dyrk1a* inhibition increases DCX^+^ neuroblast output relative to *Plagl2* alone, implicating *Dyrk1a* inhibition in the control of neurogenic fate.^23^ Given that endogenous *Dyrk1a* expression was minimal in MG, these observations suggest that *Plagl2* is sufficient to establish proliferative competence in these cells, whereas extrinsic injury-associated cues may functionally substitute for *Dyrk1a* inhibition to establish a permissive state for neurogenic competence.

The establishment of this permissive state was likely facilitated by the specific architecture of our genetic framework. For instance, the use of the *Hes5* promoter offers a distinct advantage over commonly used flox-based paradigms that enforce constitutive transgene overexpression or target gene deletion.^11,12,14,17,35^ Unlike these static systems, the *Hes5*-driven iPaD transgene remains active during MG reprogramming, but is naturally reduced with neuronal maturation, which may better facilitate the transition from proliferation to neuronal fate commitment. However, in the context of AAV-mediated reprogramming, the shortened *Hes5* promoter showed endogenous leakage into neurons, similar to that reported for *Gfap* minipromotor-based AAV systems.^41^ Notably, this leakage appeared more prominent in *AAV-Plagl2* retinas, raising the possibility that the transgene itself may enhance promoter activity through cis-regulatory effects. Nevertheless, the transgenic and AAV-based systems both supported robust *Plagl2*-dependent MG proliferation, but neurogenic output was markedly reduced in the AAV setting, even following NMDA injury. This difference may reflect the progressive dilution of AAV-driven *Plagl2* expression during repeated rounds of cell division, such that *Plagl2* expression becomes insufficient to support neuronal differentiation.

At the transcriptional level, Wnt signaling and *Ccnd1* were upregulated in *Plagl2*-activated MG, both of which have been implicated as downstream targets of *Plagl2* and as drivers of MG proliferation.^10,13,23,37^ Notably, neurogenic genes that are targeted by *Plagl2* in NSCs, such as *Ascl1* and *Neurog2*, were similarly upregulated in *Plagl2*-activated MG regardless of injury status, supporting the conservation of core *Plagl2*-driven transcriptional programs across cell lineages. Consistent with this, *Plagl2*-activated MG following NMDA injury showed an increased fraction of the neurogenic MG cluster, which most closely resembled a late RPC-like transcriptional state. This shift toward an RPC-like state may reflect the temporal dynamics of induced neurogenic factors. In aged hippocampal NSCs, iPaD induces endogenous ASCL1 oscillations, a feature that is functionally important because *Ascl1* exhibits distinct dynamics-dependent functions in NSCs, i.e., sustained expression promotes neuronal fate commitment, whereas oscillatory expression maintains multipotency and proliferation.^25,30^ We found that *Plagl2*-activated MG exhibited endogenous ASCL1 oscillations by 72 hpi, a stage at which mammalian MG would be expected to return toward a resting state.^5,12^ Notably, the periodicity of ASCL1 oscillations resembled that observed in active NSCs and RPCs. This distinguishes *Plagl2*-driven reprogramming from prior paradigms that relied on sustained exogenous ASCL1 expression together with NMDA injury and chromatin remodeling, which favored direct transdifferentiation with limited proliferative expansion.^14,35^ We speculate that this outcome is a direct consequence of sustained *Ascl1* expression, which may drive premature neuronal fate commitment by bypassing an intermediate proliferative state. Together, these results support a model in which *Plagl2* is sufficient to initiate MG proliferation, whereas injury-associated cues act synergistically to drive endogenous ASCL1 oscillations, thereby stabilizing a proliferative yet neurogenically primed state that closely resembles late RPCs. More broadly, such dynamics-dependent transcription factor control is not unique to *Ascl1* or to the central nervous system and has been described across multiple developmental systems.^45–49^ A key remaining question is how cells decode oscillatory versus sustained inputs to direct distinct lineage outcomes.

At the level of neuronal output, immunohistochemical and scRNA-seq analyses revealed that *Plagl2*-driven MG reprogramming remains constrained to bipolar/amacrine fates, consistent with the developmental competence of late RPCs to generate later-born retinal neurons and prior studies in which bipolar neurons are among the most frequent MG-derived neurogenic outputs.^14–17,32^ Nonetheless, this fate bias highlights an important translational limitation, as PRs and RGCs are among the most clinically relevant retinal neuronal subtypes for regenerative repair.^50^ This limitation persists despite clear evidence of migratory plasticity of *Plagl2*-activated MG toward the PR-resident ONL and the RGC-resident GCL. Previous studies indicate that co-expression of additional bHLH factors or developmentally active transcription factors can potentiate MG-to-neuron conversion efficiency and expand lineage output toward RGC-like states, but the induced cells largely remain in the INL.^15,35^ In contrast, *Plagl2*-activated MG migrated toward the GCL following NMDA injury, yet failed to acquire RGC-like transcriptomic features, indicating that positional migration and terminal fate specification can be uncoupled. Despite this, multielectrode array recordings showed only the partial recovery of light-evoked RGC output following *Plagl2* induction. This suggests that functional improvement is more likely mediated indirectly through the enhanced generation of INL neurons, which may integrate into existing retinal circuitry and improve signal propagation to RGCs, rather than through the direct replacement of RGCs themselves. Similarly, despite the late RPC-like bias and the fact that PRs are the dominant late-born retinal neurons, ONL-resident *Plagl2*-activated MG did not acquire PR-like morphology, suggesting that late RPC-like competence alone is insufficient to explain PR fate output in this context and that additional barriers to reprogramming competence remain. Consistently, although the iPaD transgene was efficiently activated in most SOX9^+^ MG, a subset of mCherry^+^ MG retained glia-like morphology in the INL without BrdU incorporation. Likewise, scRNA-seq showed that a subset of *Plagl2*-activated MG remained in the MG cluster, which was enriched for glial-associated genes, regardless of injury status. Together, these findings suggest that efficient MG reprogramming will require not only induction of pro-neurogenic programs, but also the concurrent suppression of pro-glial state regulators to overcome barriers to lineage diversification toward clinically relevant retinal subtypes.

Despite these limitations, the ability of *Plagl2* to unlock the latent regenerative potential of MG supports a broader conceptual shift in how we approach tissue repair. The inter-tissue activity of *Plagl2* parallels other potent reprogramming paradigms that enhance regenerative plasticity across diverse tissues and disease contexts. A prominent example is *in vivo* partial reprogramming with Yamanaka factors, which alleviates pancreas and muscle injury,^51^ improves RGC axonal regeneration,^52^ and attenuates the molecular hallmarks of Alzheimer’s disease.^53^ In the context of MG, AAV-mediated *Oct4* overexpression reportedly promotes MG transdifferentiation toward a bipolar fate.^54^ Alternatively, *Ptbp1* knockdown has also been proposed as a transferable glia-to-neuron conversion paradigm across multiple regions of the central nervous system, although the extent of MG-to-neuron conversion remains actively debated.^55–57^ Together, these studies point to an alternative to existing inter-species-inspired MG reprogramming strategies, in which regenerative competence is accessed by redeploying reprogramming paradigms identified in other mammalian lineages, positioning *Plagl2* as a novel transferable regulator of regenerative competence across distinct cell lineages.

## Limitation of the study

While previous studies have commonly employed genetic lineage tracing (i.e. *GlastCreER;Sun1-GFP*) to definitively track the fate of MG and MG-derived progeny following reprogramming, our study utilized the *Hes5* promoter to drive transgene expression. Given that *Hes5* activity diminishes as cells undergo neuronal maturation, this approach may underestimate the full extent of *Plagl2*-induced reprogramming. Another limitation is that, although multielectrode array recordings suggest partial recovery of light-evoked RGC output, the mechanism underlying this improvement remains unclear. In particular, it is not yet possible to determine whether MG-derived INL neurons indirectly enhance RGC response through integration into upstream retinal circuitry, or whether *Plagl2*-activated MG in the GCL contribute more directly to the measured output.

## Acknowledgements

We thank members of the Kageyama lab and Masayo Takahashi for discussions, Kotari Ishikawa for assisting mice experiments, as well as Tom Reh and Stefanie Wohl for guidance with the dissociation protocol. We also thank the staff of Research Resources Division, RIKEN, Center for Brain Sciences, for assistance with FACS analysis and animal maintenance. This work was supported by: Grant-in-Aid for Scientific Research from Japan Society for the Promotion of Science (JSPS) to T.M. (24KJ1350), and Specially Promoted Research from JSPS to R.K. (21H04976).

## Author contributions

T.M. performed the experiments, analyzed the data, and wrote the manuscript; M.W. and M.M. conducted multielectrode array experiments and analysis; M.S. and T.A. generated *Achilles-Ascl1* knock-in mice; R.K. supervised the project and wrote the manuscript. All authors reviewed and approved the final manuscript.

## Methods

### Resource availability

Further information and requests for resources and reagents should be directed to and will be fulfilled by the lead contact, Ryoichiro Kageyama

### Materials availability

Newly generated items in the study are available on request from the lead contact with a completed materials transfer agreement.

### Data and code availability

- The scRNA-seq data are available in the DDBJ Sequence Read Archive (DRA) under BioProject PRJDB40589 (submission PSUB047818), and genomic expression archive (GEA) with accession number: E-GEAD-1226 (submission ESUB002561).
- This paper does not report original code
- Any additional information needed to reanalyze the data reported in this paper is available from the lead contact upon request.

### Experimental model and subject details

#### Animals

Animal experiments were approved by the Animal Experiment Committee of RIKEN. Mice were housed in polysulfone cages within a specific-pathogen-free facility under a 12-hour light/dark cycle, with *ad libitum* access to water and standard chow (Oriental Yeast). *pHes5-mCherry* and *pHes5-iPaD-mCherry* transgenic mice were established previously on an Institute of Cancer Research (ICR) background (JSLC).^25,29^ *Achilles-Ascl1* knock-in mice (Accession No. CDB0374E: https://large.riken.jp/distribution/mutant-list.html) were generated on a C57BL/6N background (CLEA Japan) using CRISPR/Cas9-mediated knock-in in zygotes to insert Achilles cDNA in-frame into the 5′ region of the *Ascl1* locus, as described previously.^58^ The gRNA site (5’-TTG CCA GAG CTC TCC ATG CC-3’) was designed using CRISPRdirect.^59^ For the homologous recombination-mediated knock-in, the donor vector consisting of homology arms and Achilles was generated to insert Achilles at 3 bases downstream of the PAM sequence. Genotyping was performed by PCR using primers, 5′-GAGGCTCCCGAAGCCAACC-3′ and 5′-GCTCCACTCTCCATCTTGCCAGA-3′ (Fig. S3C). Homozygous animals (∼2 months old) of both sexes were used for *pHes5-iPaD-mCherry* mice and knock-in (*Achilles-Ascl1*) lines, whereas homozygous and heterozygous animals of both sexes were used for the *pHes5-mCherry* line.

### Method details Doxycycline

#### treatment

Doxycycline (1 mg/mL; Sigma) was dissolved in drinking water and administered *ad libitum* to adult mice for the indicated period.

### BrdU/EdU treatment

BrdU (1 mg/mL; Sigma) or EdU (0.5 mg/mL; Fujifilm Wako) was dissolved in drinking water and administered *ad libitum* to adult mice for the indicated period.

### NMDA injection

Adult mice were anesthetized with a triple-anesthetic mixture consisting of Domitor (0.75 mg/kg; Zenoaq), midazolam (4 mg/kg; Sandoz), and butorphanol (5 mg/kg; Meiji), administered intraperitoneally at a volume of 0.1 mL per 10 g body weight. 2 µL of NMDA (100 mM; Selleck), dissolved in D-PBS(–) (Nacalai) and supplemented with 0.05% Fast Green (Fujifilm Wako), was injected intravitreally into the right eye. A small puncture was first made with a 33-gauge needle (Nipro), after which 2 µL of NMDA solution was injected using a Hamilton syringe fitted with a pulled, blunt-end glass capillary. After the procedure, atipamezole (0.75 mg/kg; Zenoaq) was administered intraperitoneally at 0.1 mL per 10 g body weight, and mice were placed on a heating pad until recovery.

### AAV production and injection

The AAV2.7m8 capsid (No. 64839) and pAdDeltaF6 helper (No. 112867) plasmids were obtained from Addgene. To enhance transgene expression, two point mutations converting surface tyrosine residues to phenylalanine (Y444F and Y730F) were introduced into the AAV2.7m8 capsid plasmid using the PrimeSTAR Mutagenesis Basal Kit (Takara). AAV constructs were transfected into the AAVpro^®^ 293T Cell Line (Takara), and viral particles were purified using the AAVpro^®^ Purification Kit (Takara). For AAV delivery, adult mice were anesthetized as described for NMDA injection, and 2 µL of the purified AAV solution was injected intravitreally into both eyes.

### Immunohistochemistry

Adult mouse eye globes were fixed in 4% paraformaldehyde (Sigma) for 4 hours at 4°C. Retinas were dissected in PBS and incubated in 30% sucrose (Nacalai) overnight at 4°C. The tissues were then embedded in O.C.T. compound (Sakura Finetek) and frozen at –80°C. Frozen retinas were sectioned at a thickness of 25 µm on a cryostat (Leica) and stored at –80°C until staining. For immunohistochemistry, sections were air-dried for 30 min at 37°C and washed for 10 min in PBS. Antigen retrieval was performed in citrate buffer (pH 6.0) for 5 min at 90°C. For BrdU detection, sections were incubated in 2 N HCl (Nacalai) for 30 min at 37°C, followed by neutralization in 0.1 M sodium tetraborate decahydrate (Nacalai) for 10 min at room temperature. For EdU detection, the Click-iT EdU Cell Proliferation Kit (Invitrogen) was used according to the manufacturer’s instructions. Sections were then incubated in blocking solution [5% donkey serum (Jackson ImmunoResearch) in 0.5% PBST] for 2 hours at room temperature, followed by incubation with primary antibodies diluted in blocking solution overnight at 4°C. After washing three times for 5 min in PBST to remove excess primary antibodies, sections were incubated with secondary antibodies in blocking solution for 3 hours at room temperature. Hoechst 33342 (1:1000; ThermoFisher) was added with secondary antibodies for nuclear counterstaining. Finally, sections were washed three times for 5 min in PBST, mounted with Fluoromount-G (Invitrogen) under coverslips, and stored at 4°C until imaging. The primary antibodies used were as follows: goat anti-mCherry (1:500; SICGEN), rat anti-BrdU (1:500; Abcam), rabbit anti-SOX9 (1:500; Sigma), rabbit anti-ASCL1 (1:250; Abcam), rabbit anti-PLAGL2 (1:250; Proteintech), rabbit anti-DYRK1A (1:250; CST), goat anti-hOTX2 (15 µg/mL; R&D Systems), rabbit anti-NEUN (1:250; Proteintech), mouse anti-HUC/D (15 µg/mL; Invitrogen), rabbit anti-BRN3A (1:250; Abcam), and rabbit anti-Recoverin (1:250; Proteintech). The secondary antibodies used were as follows: Alexa Fluor 405-conjugated anti-rabbit IgG (1:500; ThermoFisher), Alexa Fluor Plus 405-conjugated anti-rabbit IgG (1:500; ThermoFisher), Alexa Fluor 488-conjugated anti-rabbit IgG (1:500; ThermoFisher), Alexa Fluor 488-conjugated anti-rat IgG (1:500; ThermoFisher), Alexa Fluor 594-conjugated anti-goat IgG (1:500; ThermoFisher), and Alexa Fluor 647-conjugated anti-rat IgG (1:500; ThermoFisher).

### Confocal fluorescent imaging and time-lapse imaging

Fluorescent images were captured using a Zeiss LSM 980 inverted confocal microscope. Figure images were acquired with a Plan-Apochromat 63x/1.4 Oil DIC objective lens using the Airyscan2 detector. For cell quantification, images were acquired with a Plan-Apochromat 20x/0.8 objective lens at 1.5 µm z-intervals. Fluorescent time-lapse imaging was performed using a Zeiss LSM 980 inverted confocal microscope with a Plan-Apochromat 20x/0.8 objective, equipped with an environmental chamber (Tokai Hit) maintained at 37°C with 5% CO_2_ and 40% O_2_. For adult retinal slice imaging, retinas were dissected in ice-cold HBSS(–) (Gibco) and manually sliced using a surgical knife. The slices were embedded in Type I-A collagen (Fujifilm Wako) prepared according to the manufacturer’s instructions and placed on a φ27-mm glass-bottom dish (Iwaki). The slices were then covered with 2 mL of culture medium consisting of 5% fetal bovine serum (FBS; Nichirei Biosciences), 5% horse serum (Gibco), 1x B27 (Gibco), 1x N-2 MAX Media Supplement (R&D Systems), 10 ng/mL EGF (ThermoFisher), 10 ng/mL bFGF (Fujifilm Wako), 100 U/mL penicillin (Nacalai), and 100 µg/mL streptomycin (Nacalai) in DMEM/F-12 (Gibco). Achilles and mCherry fluorescence were excited at 514 nm and 561 nm, respectively. For RPC imaging, retinas from *Achilles-Ascl1;pHes5-iPaD-mCherry* embryos (E13.5) were dissected in ice-cold HBSS(–) (Gibco), and individual eyes were dissociated using the Papain Dissociation System (Worthington), prepared according to the manufacturer’s instructions. Retinas were incubated in papain/DNase solution for 10 min at 37°C, then centrifuged for 5 min at 300 × *g* at 4°C. The resulting cell pellets were resuspended in DMEM/F-12 (Gibco) and washed three times. Dissociated cells were subsequently plated in a 35-mm dish coated with Matrigel^®^ (1:400, Corning) and the culture medium described above without serum. Cells were passaged at least three times prior to imaging.

### Single cell tracking and analysis

Time-lapse imaging data were processed in Fiji using a Gaussian filter, and single-cell tracking was manually performed with Mastodon. Subsequent analyses were conducted in R. Briefly, Achilles and mCherry fluorescence intensities were normalized within each track and smoothed using a Savitzky-Golay filter. To determine signal periodicity, detrending was performed on the normalized intensities by subtracting moving-average values. Periodic oscillations were then analyzed by wavelet transform using the analyze.wavelet function.

### Multielectrode array recordings and analysis

RGC activity was recorded using a multielectrode array system (Multi Channel Systems, 60pMEA200/30iR-Ti), as previously described with minor modifications.^34^ Mice were dark-adapted for at least 1 hour before the experiments. After enucleation, the retina was isolated and placed RGC-side down onto the multielectrode array. The tissue was fixed by gentle suction and continuously perfused with Ames’ medium (United States Biological, A1372-25) at 34°C with 95% O_2_ and 5% CO_2_ at 3–3.5 mL/min. Extracellular signals were amplified using an multielectrode array amplifier (Multi Channel Systems, USB-ME64-System) and digitized at 20 kHz. For analysis, units were included if: waveform amplitude was ≥ 20 µV, and mean firing rate ≥ 0.1 Hz. PSTHs were constructed for each unit based on raster plots obtained from three trials of flash stimulation using 20-ms bins. Spike-sorted RGC units were analyzed for flash-evoked responses. Baseline firing rate was calculated from spontaneous activity during the 5 seconds prior to stimulus onset. The ON response window was defined as 0 to 0.5 s, and the OFF response window was defined as 1.05 to 1.35 s relative to flash onset. Peak firing rates were calculated within each response window after subtracting baseline activity. For each polarity (ON or OFF), a response was considered positive only when all of the following criteria were satisfied: (1) the baseline-corrected peak firing rate exceeded the baseline-derived threshold (defined as 4 × SD of baseline activity), (2) at least one spike was detected in the corresponding response window in at least two stimulus trials, and (3) the total number of spikes within the response window across trials was at least three. To enable statistical comparison of peak firing rates between groups, data were summarized at the mouse level. For each retina, peak firing rates were summarized as the median across included units. When both eyes were recorded, values were averaged to obtain a single animal-level metric.

### Light stimulation

Full-field light stimulation was delivered using a 505 nm LED (Thorlabs, SOLIS-505C) controlled by an LED driver (Thorlabs, DC2200) and a function generator (NF Corporation, WF1973). Flash stimulation consisted of 1-second light pulses at an intensity of 14,189 R*/rod/s under dark-adapted conditions.

### Single-cell dissociation for GEM-X library preparation

Retinas were dissected in ice-cold HBSS(–) (Gibco) and then dissociated using the Papain Dissociation System (Worthington). Each sample comprised a minimum of 6 retinas from animals of both sexes. Retinas were dissociated in papain/DNase solution for 10 min at 37°C, after which ovomucoid/DNase solution was added to terminate papain activity. The suspension was gently triturated using a P1000 pipette until it became uniformly cloudy. Dissociated cells were centrifuged for 10 min at 300 × *g* at 4°C, resuspended in ice-cold HBSS(+) containing 5% FBS (Nichirei Biosciences), 0.05% BSA (Gibco), 100 U/mL DNase (Worthington), and 4′,6-diamidino-2-phenylindole (DAPI; 1:1000; Invitrogen) to label dead cells, and filtered through a 35 µm cell strainer (Falcon) for FACS. FACS was performed on a BD FACSAria III Cell Sorter (BD Biosciences), and the mCherry-positive, DAPI-negative fraction was collected. The sorted cells were cryopreserved (∼2000 cells/µL) in Stem-CellBanker^®^ (Takara) and sent to Novogene for GEM-X library preparation and sequencing.

### Single-cell RNA sequencing analysis

Sequencing results were preprocessed and aligned to the GRCm39 genome with 10x Genomics Cell Ranger (v9.0.1). Downstream analyses were performed in R (v4.5.2) using Seurat (v5.4.0).^60^ For each sample, cells with < 4,000 or > 30,000 detected UMIs, or with > 10% mitochondrial gene expression, were removed. Gene expression values were normalized using log normalization, and 3,000 highly variable genes were identified using the variance-stabilizing transformation method. Principal component analysis was performed on the variable genes, and clusters displaying low UMI counts, or low cell number were further removed. Filtered datasets were integrated using Harmony (v1.2.4),^61^ with sample identity specified as the batch variable, and UMAP dimensional reduction was applied on the first 20 harmony-corrected embeddings. Cluster-specific marker genes were identified using FindAllMarkers function with the Wilcoxon rank-sum test. Gene ontology analysis was performed using the clusterProfiler package.^62^ For comparison with the developing retina, a previously annotated reference dataset was used.^32^ Transfer anchors between the reference and query datasets were identified using the FindTransferAnchors function, and reference cell-type annotations were transferred to the query dataset using the MapQuery function. For pseudotime analysis, MG, rMG, and NeuroMG clusters were analyzed using Palantir.^63^ Gene modules were determined with Monocle3’s find_gene_modules function and ordered along pseudotime.^64^

### Quantification and statistical analysis

#### Statistical analysis

A minimum of three fields per retina were imaged and averaged for each sample. Statistical analyses and data visualization were performed in R. Data are presented as mean ± SEM in all figures. Sample sizes are indicated in the corresponding figure legends. Two-group comparisons were performed using an unpaired Student’s *t*-test, and multi-factor data were analyzed by ANOVA followed by post hoc pairwise comparisons with Tukey’s test. Comparisons not indicated were not significant. n.s., p > 0.05; *p < 0.05; **p < 0.01; ***p < 0.001.

## Supplemental Data

### Supplemental Tables

**Table S1:** Differentially expressed genes identified from annotated scRNA-seq clusters.

**Table S2:** Shared genes between PLAGL2 ChIP-seq targets in NSCs and genes upregulated in MG.

### Supplemental Movies

**Movie S1:** *Plagl2*-activated MG undergoing multiple rounds of cell division.

**Movie S2:** *Plagl2*-activated MG exhibiting oscillatory ASCL1 expression.

---

**Figure S1.**
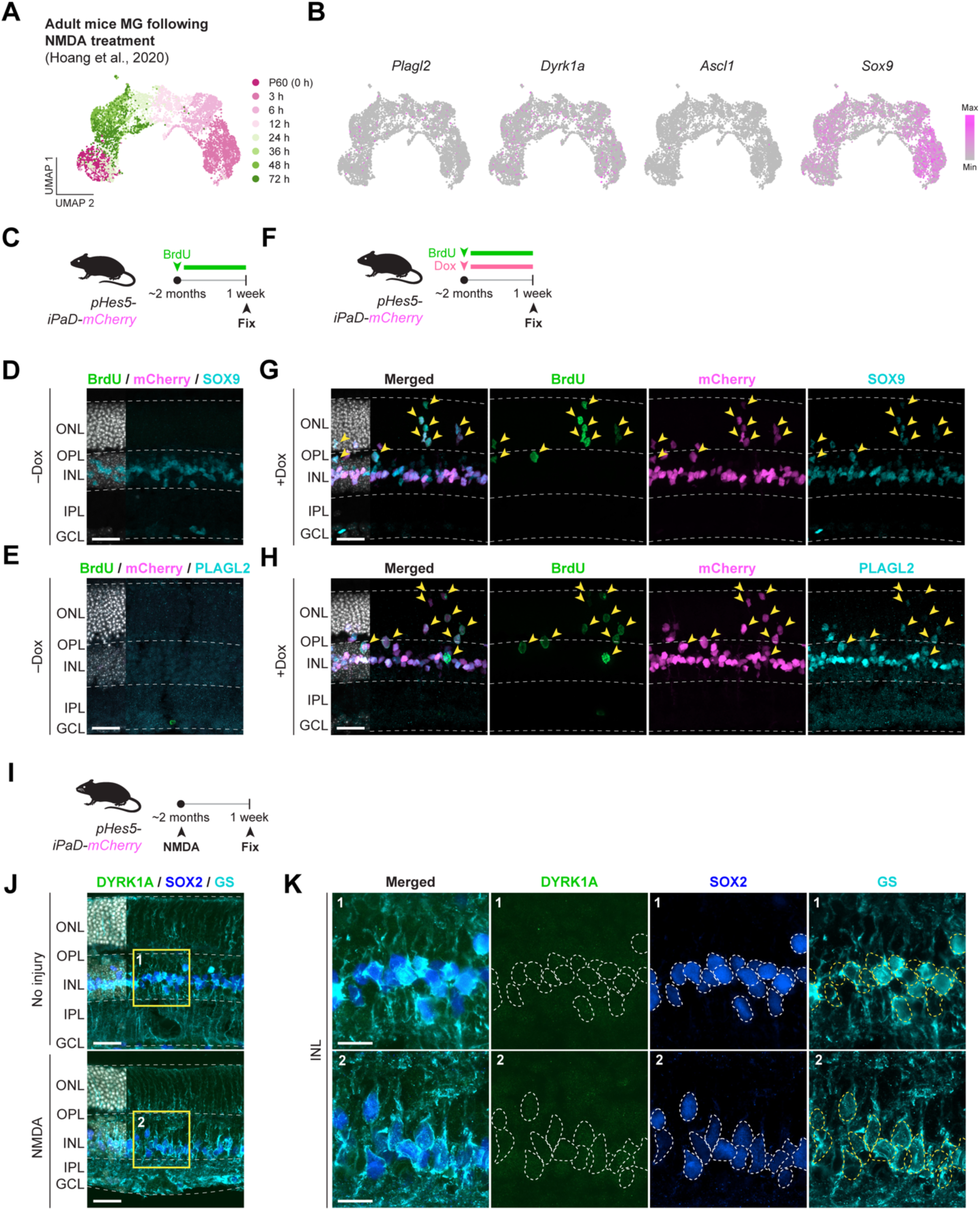
*Plagl2* activates MG proliferation without NMDA injury. A) Integrated UMAP visualization of time-course scRNA-seq data from adult mouse retinal cells following NMDA injury. B) Expression of *Plagl2*, *Dyrk1a*, *Ascl1*, and *Sox9* expression in adult mouse MG retina following NMDA injection. C) Experimental paradigm used for negative control without Dox treatment. D) Representative confocal images of *pHes5-iPaD-mCherry* retinas shown in (C), immunostained for BrdU (green), mCherry (magenta), and SOX9 (cyan). E) Representative confocal images of *pHes5-iPaD-mCherry* retinas shown in (C), immunostained for BrdU (green), mCherry (magenta), and PLAGL2 (cyan). F) Experimental paradigm used to activate iPaD transgene. G) Representative confocal images of *pHes5-iPaD-mCherry* retinas shown in (F), immunostained for BrdU (green), mCherry (magenta), and SOX9 (cyan). Yellow arrowheads represent BrdU^+^/mCherry^+^ cells. Scale bar, 25 µm. H) Representative confocal images of *pHes5-iPaD-mCherry* retinas shown in (F), immunostained for BrdU (green), mCherry (magenta), and PLAGL2 (cyan). Yellow arrowheads represent BrdU^+^/mCherry^+^ cells. Scale bar, 25 µm I) Experimental paradigm used to assess endogenous DYRK1A expression. J) Representative confocal images of *pHes5-iPaD-mCherry* retinas shown in (I), immunostained for DYRK1A (green), SOX2 (blue), and glutamine synthetase (GS; cyan). Scale bar, 25 µm. K) Magnified images from (J). Scale bar, 10 µm.

**Figure S2.**
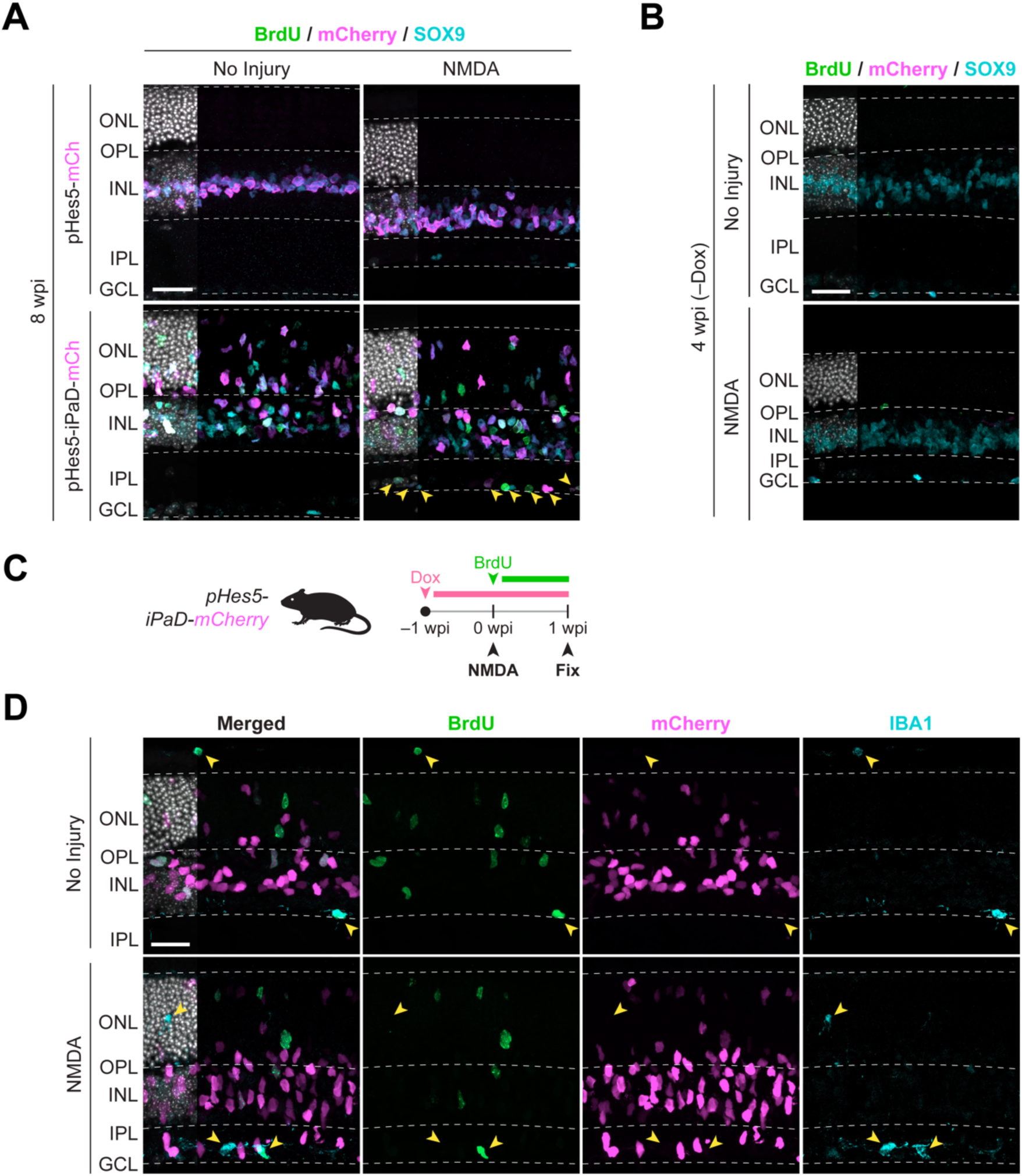
*Plagl2* activates MG proliferation without NMDA injury. A) Representative confocal images of *pHes5-mCherry* and *pHes5-iPaD-mCherry* retinas at 8 wpi, immunostained for BrdU (green), mCherry (magenta), and SOX9 (cyan). Yellow arrowheads indicate mCherry^+^ cells that migrated toward the GCL. Scale bars, 25 µm. B) Representative confocal images of *pHes5-iPaD-mCherry* retinas without doxycycline treatment at 4 wpi, immunostained for BrdU (green), mCherry (magenta), and SOX9 (cyan). Scale bar, 25 µm. C) Experimental paradigm used to activate iPaD transgene. D) Representative confocal images of *pHes5-iPaD-mCherry* mice shown in (C), immunostained for BrdU (green), mCherry (magenta), and IBA1 (cyan). Yellow arrowheads represent IBA1^+^ microglia. Scale bar, 25 µm.

**Figure S3.**
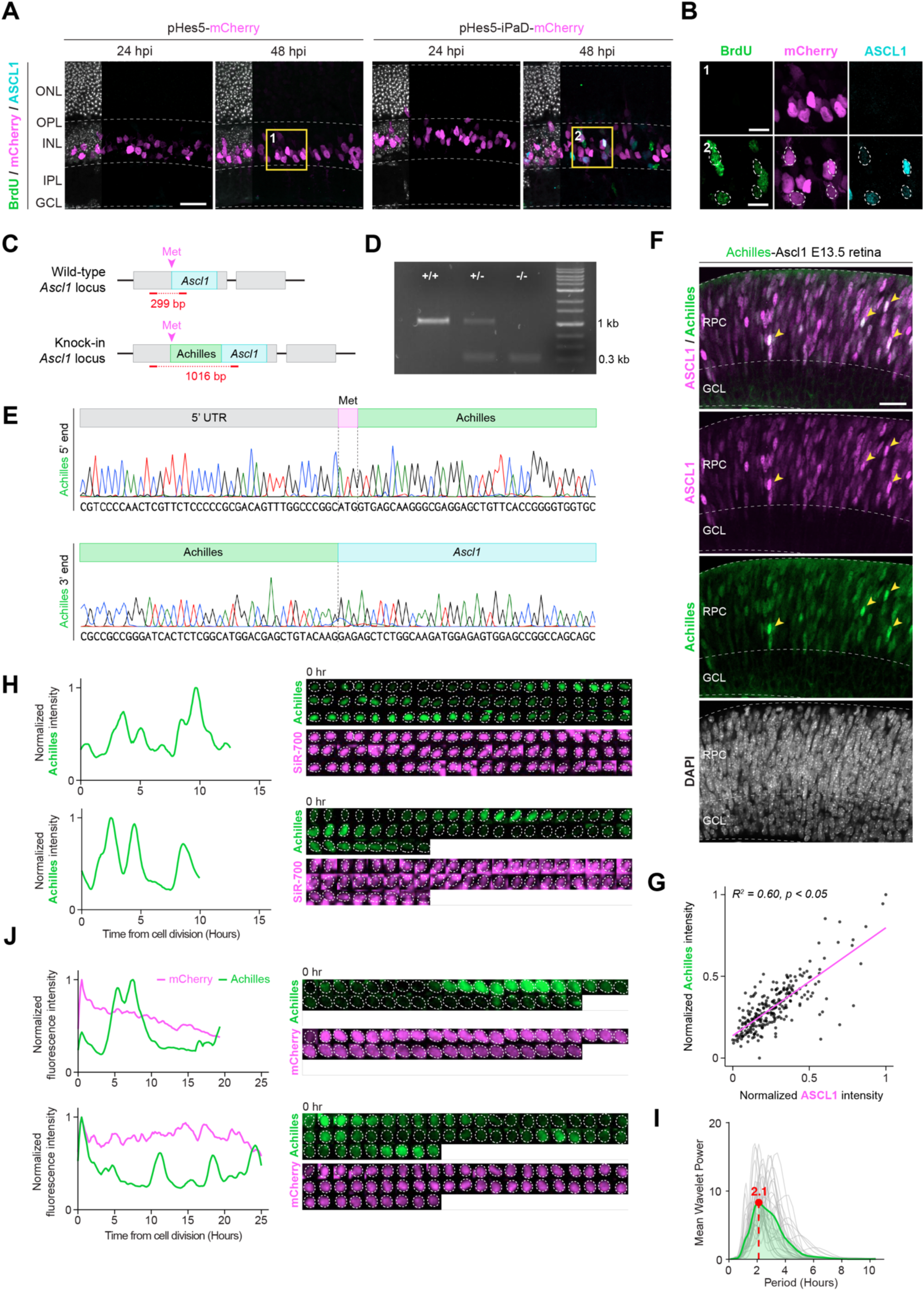
*Plagl2* induces ultradian ASCL1 oscillations in MG following NMDA injury. A) Representative confocal images of *pHes5-mCherry* and *pHes5-iPaD-mCherry* retinas at 24 and 48 hpi, immunostained for BrdU (green), mCherry (magenta), and ASCL1 (cyan). Scale bars, 25 µm. *n* = 3–4 retinas per group. B) Magnified images of ASCL1^+^/mCherry^+^ cells denoted by dotted lines from (A). Scale bar, 10 µm. C) Schematic of the wild-type and Achilles knock-in *Ascl1* loci. D) Genotyping of *Achilles-Ascl1* knock-in homozygous, heterozygous, and wild-type mice using the primers shown in (C). E) An example of sequencing confirming the correct integration of Achilles in-frame at the 5’ end of the *Ascl1* locus. F) Representative confocal images of *Achilles-Ascl1* embryonic retina, immunostained for Achilles (green) and endogenous ASCL1 (magenta). Yellow arrowheads indicate cells with high Achilles and ASCL1 co-expression. Scale bar, 25 µm. G) Quantification of normalized Achilles and endogenous ASCL1 fluorescence intensities. H) Representative single-cell tracks of *Achilles-Ascl1* fluorescence intensity traces from primary cultured RPCs derived from *pHes5-iPaD-mCherry;Achilles-Ascl1* mice. I) Mean wavelet power spectra for Achilles fluorescence signals of RPCs shown in (H). *n* = 30 tracks. J) Additional single-cell tracks from time-lapse imaging of *Plagl2*-activated MG showing *Achilles-Ascl1* and *iPaD-mCherry* fluorescence intensities.

**Figure S4.**
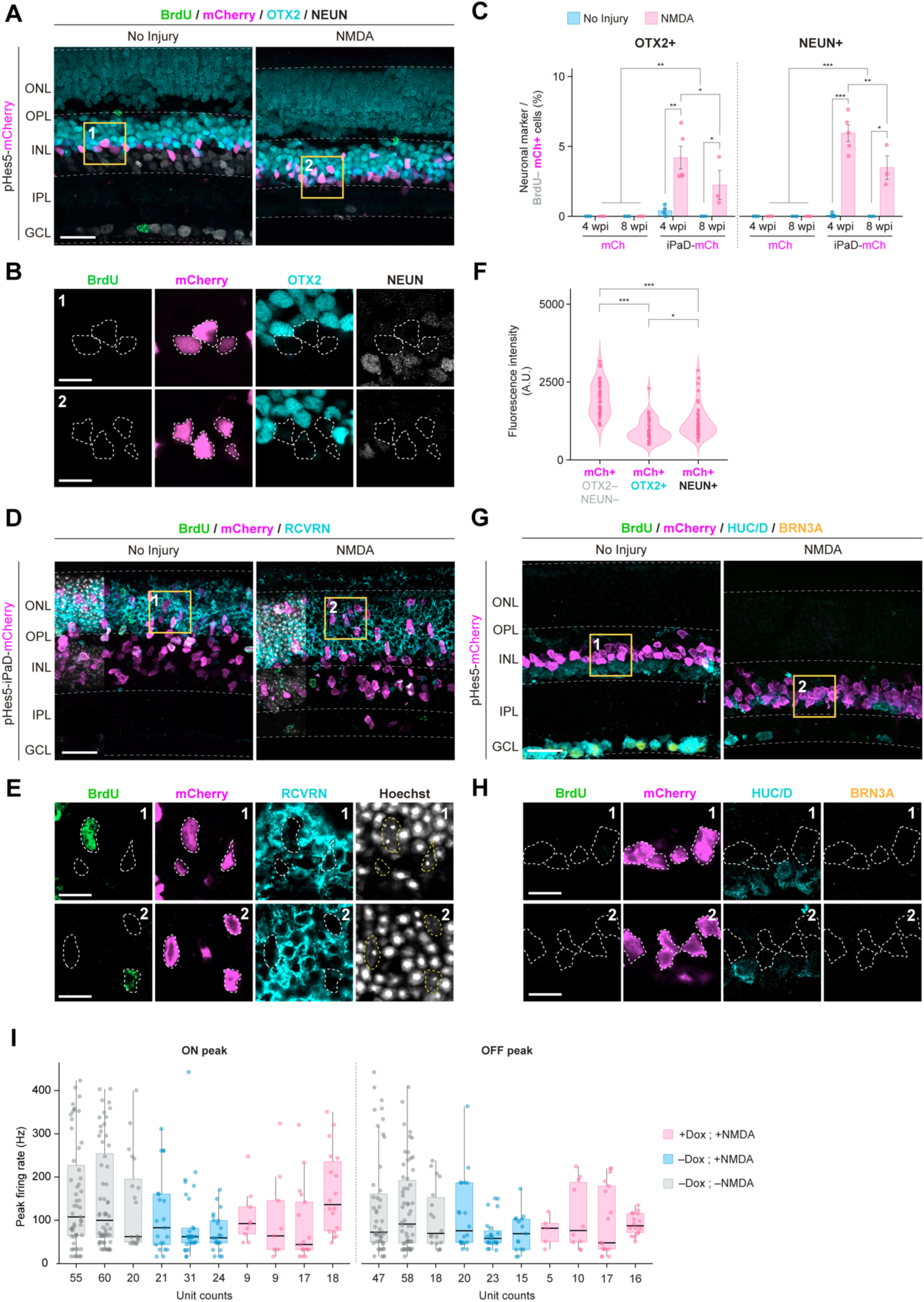
*Plagl2*-activated MG initiate neurogenesis toward INL neurons upon NMDA injury. A) Representative confocal images of control *pHes5-mCherry* retinas at 4 wpi with or without NMDA injury. Sections were immunostained for BrdU (green), OTX2 (cyan), and NEUN (white). mCherry (magenta) signal was detected by native fluorescence. Scale bars, 25 µm. B) Magnified images of mCherry^+^ cells from (A). Scale bar, 10 µm. C) Proportion<u>s</u> of BrdU^−^/mCherry^+^ cells co-expressing OTX2 or NEUN. Data are presented as mean ± SEM. Significance was determined by three-way ANOVA followed by post hoc Tukey’s test. *p < 0.05, **p < 0.01, ***p < 0.001. *n* = 3–5 retinas per group. D) Representative confocal images of control *pHes5-mCherry* retinas at 4 wpi with or without NMDA injury. Sections were immunostained for BrdU (green), mCherry (magenta), and recoverin (cyan). Scale bars, 25 µm. E) Magnified images of mCherry^+^ cells from (D). Scale bar, 10 µm. F) Fluorescence intensity of mCherry^+^ cells in *pHes5-iPaD-mCherry* retinas with NMDA injury classified as OTX2^−^/NEUN^−^, OTX2^+^, or NEUN^+^. Each dot represents a single cell. Significance was determined by one-way ANOVA followed by post hoc Tukey’s test. *p < 0.05, ***p < 0.001. *n* = 40 cells per group from 4 retinas. G) Representative confocal images of control *pHes5-mCherry* retinas at 4 wpi with or without NMDA injury. Sections were immunostained for BrdU (green), mCherry (magenta), HUC/D (cyan), and BRN3A (yellow). Scale bars, 25 µm. H) Magnified images of mCherry^+^ cells from (G). Scale bar, 10 µm. I) Mouse-level distributions of corrected ON and OFF peak firing rates across experimental conditions. Each dot represents one RGC unit and each black line represents the mouse-level summary value (each dot in Fig. 4J), calculated as the mean across eyes of the eye-level median unit peak. The ON peak plot includes ON and ON-OFF units, and the OFF peak plot includes OFF and ON-OFF units. *n* = 3–4 mice per group.

**Figure S5.**
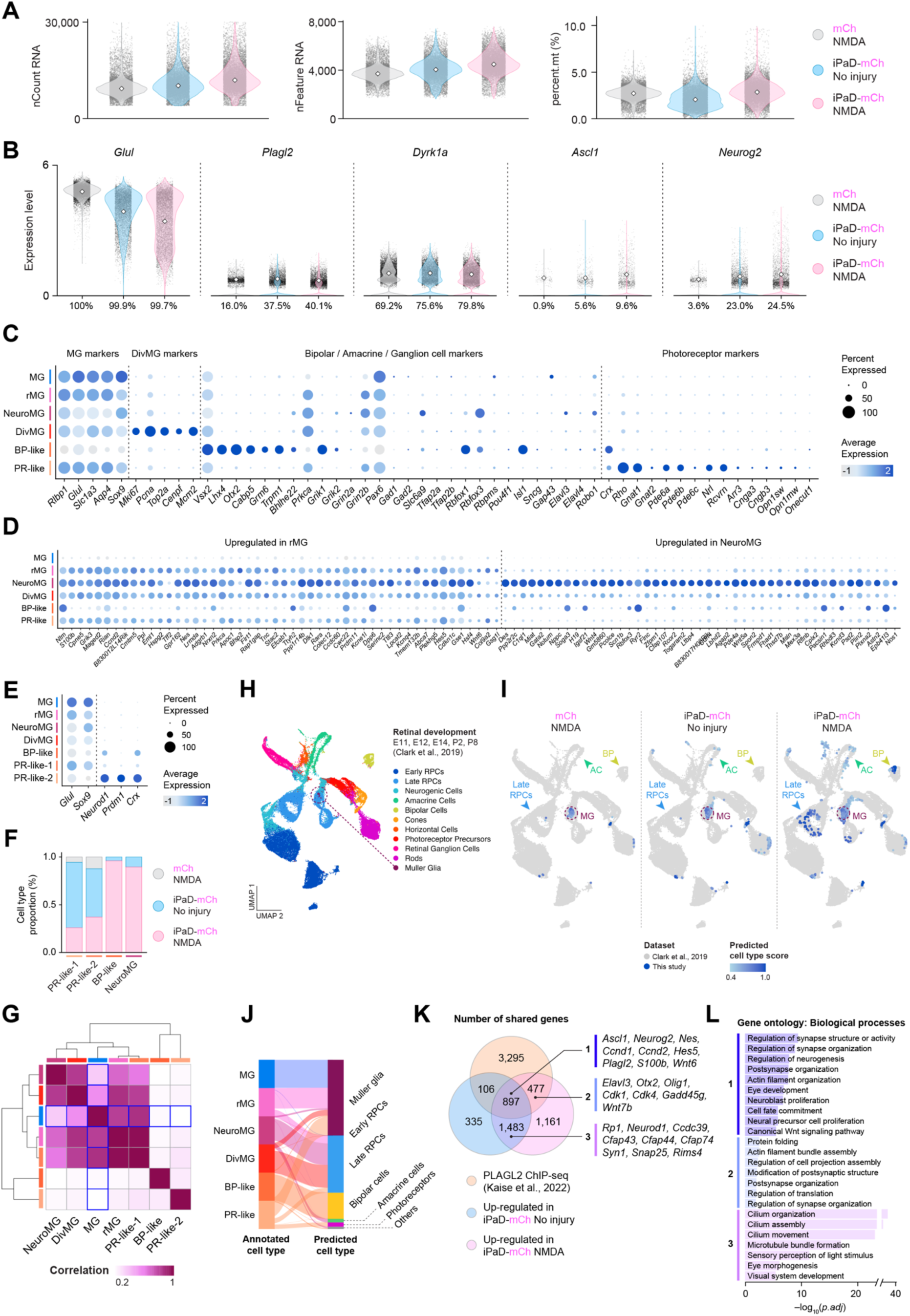
scRNA-seq reveals progressive neurogenic reprogramming of MG following *Plagl2* induction. A) Quality-control metrics for scRNA-seq dataset. B) Violin plots showing *Glul*, *Plagl2*, *Dyrk1a*, *Ascl1*, and *Neurog2* expression across each condition. Percent values represent fraction of cells with detectable expression. White squares denote mean expression among expressing cells. C) Dot plot of canonical MG markers, cell cycle genes, and mature retinal neuronal markers across annotated clusters. D) Dot plot of genes upregulated in reprogrammed MG (rMG; left) and neurogenic MG (NeuroMG; right) across the annotated clusters. E) Dot plot of PR-associated markers across the PR-like clusters. F) Proportion of PR-like-1, PR-like-2, bipolar (BP)-like, and NeuroMG cells for each sample identity. G) Correlation heatmap of average gene expression profiles across the annotated clusters. H) Integrated UMAP of the mouse retinal developmental scRNA-seq reference dataset with pre-annotated cell clusters. I) Projection of query cells (this study) onto the reference developmental atlas. J) Alluvial plot linking assigned cluster annotation from this study to predicted retinal developmental cell types from the reference atlas. K) Venn diagram showing the overlap between PLAGL2 ChIP-seq target peaks in NSCs and genes upregulated in *pHes5-iPaD-mCherry* retinas with or without NMDA injury. L) GO biological process enrichment for overlapping gene sets.

**Figure S6.**
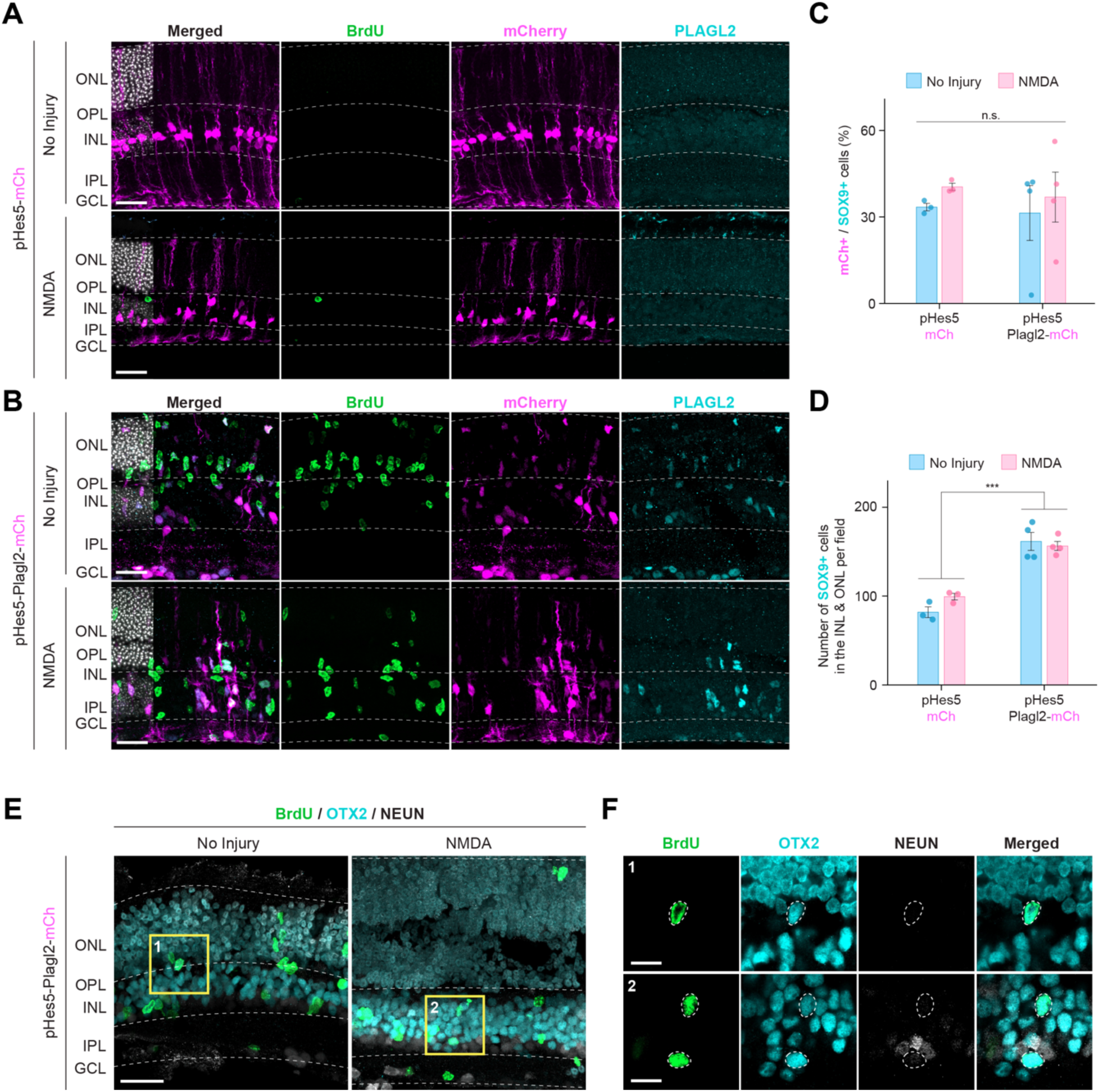
AAV-mediated *Plagl2* overexpression sufficiently induces MG proliferation. A) Representative confocal images of retinas infected with *AAV-pHes5-mCherry*, with or without NMDA injury. Sections were immunostained for BrdU (green), mCherry (magenta), and PLAGL2 (cyan). Scale bars, 25 µm. B) Representative confocal images of retinas infected with *AAV-pHes5-Plagl2-mCherry*, with or without NMDA injury. Sections were immunostained for BrdU (green), mCherry (magenta), and PLAGL2 (cyan). Scale bars, 25 µm. C) Quantification of mCherry^+^/SOX9^+^ cells in the INL and ONL. Data are presented as mean ± SEM. Significance was determined by two-way ANOVA followed by post hoc Tukey’s test. *n* = 3–4 retinas per group. D) Quantification of SOX9^+^ cells in the INL and ONL. Data are presented as mean ± SEM. Significance was determined by two-way ANOVA followed by post hoc Tukey’s test. ***p < 0.001. *n* = 3–4 retinas per group. E) Representative confocal images of retinas infected with *AAV-pHes5-Plagl2-mCherry* with or without NMDA injury. Sections were immunostained for BrdU (green), OTX2 (cyan), and NEUN (gray). Scale bars, 25 µm. F) Magnified images of OTX2^+^/BrdU^+^ cells from (E). Scale bar, 10 µm.

